# Extending cis-regulatory networks using chromatin-RNA interactions

**DOI:** 10.64898/2025.12.15.694370

**Authors:** Pelin Sahlén, Antón Carcedo, Yung-Li Chen, Samuel Kostic, Artemy Zhigulev, Madeleine Petersson Sjögren, Julie Cordier, Wing Hin Yip, Chi Wai Yip, Kayoko Yasuzawa, Takeya Kasukawa, Masaki Kato, Alice Lambolez, Hazuki Takahashi, Magda Bienko, Ludvig Lizana, Piero Carninci

## Abstract

Cis-regulatory networks are essential for determining gene transcriptional states, yet the mechanisms that mediate cellular signaling and enhancer-promoter communications are not yet well understood. Here, we integrate high-resolution enhancer-promoter interactions obtained using targeted chromosome conformation capture and RNA-DNA contacts from RADICL-seq to uncover the role of RNA in modulating enhancer-promoter interactions. Both datasets were generated from human induced pluripotent cells at different stages across neural differentiation. As expected, most enhancer-promoter interactions were dynamic with only 38% shared across all cell states and 68% of promoter interacting regions overlapped at least one annotated neural enhancer. Integration with RADICL-seq data revealed a 9.3-fold enrichment for cis-interacting DNA regions to associate with at least one RNA. Among 18,346 cis-interacting promoters, only 1,170 (6.3%) lacked any RNA association, whereas 3,702 (28%) showed stable RNA association and 13,579 (74%) displayed dynamic RNA association across differentiation. Promoters associated with RNA engaged with higher number of enhancers and exhibited higher expression levels, while enhancers with RNA associations showed higher levels of chromatin activation marks. Promoters with dynamic RNA association were also more likely to be differentially expressed and enriched for relevant biological processes, phenotypes and diseases. Importantly, RNA-DNA association dynamics correlated strongly with DNA-DNA interaction dynamics; gain of RNA association on enhancers was typically accompanied by enhancer-promoter interaction gain, whereas concurrent RNA association gain on both promoter and enhancers frequently led to interaction loss. The correlation of RNA association on cis-interaction dynamics across differentiation were also captured in the community structure of the DNA-RNA network. Together, our results reveal extensive coupling between RNA-DNA and DNA-DNA networks supporting a coordinating role of RNA in gene regulation.

## Introduction

There is a broad and functionally important class of RNAs that associate with chromatin(*1*). These chromatin-associated RNAs (caRNAs) govern genome regulation mainly in two modes: *cis* and *trans* (reviewed in (*2*)). In *cis*, nascent transcripts stay near the gene locus to help regulate transcription by modulating either polymerase pausing (*3*), transcript activation (*4*), termination (*5*) or repression (*6*) (reviewed in (*7*)). In *trans*, caRNAs diffuse or are actively transported to distant chromatin loci. Several trans-acting interactions caRNAs such as MALAT-1(*8*), NEAT1(*8*), SNHG14(*9*), HOTAIR(*10*) and Xist(*11*) have been studied in detail revealing their essential role in genome regulation. MALAT-1, for example, is a long non-coding RNA (lncRNA) that binds widely across chromatin and regulates alternative splicing by interacting with serine/arginine residues of splicing factors that alter their nuclear distribution and abundance (*12*). Similarly intergenic RNAs can localize to specific chromatin regions to maintain cellular stemness in a species-specific manner (*13*). Beyond transcriptional control, caRNAs have also architectural roles: long intergenic non-coding RNAs (lincRNAs) can scaffold RNA-binding proteins that form nuclear compartments (*14*), and caRNAs contribute to three-dimensional genome organisation by localising near topologically associating domain borders and enhancer-promoter loops (*15–18*).

While genome-wide RNA-DNA mapping methods such as GRID-seq(*16*), ChAR-seq(*19*) and RADICL-seq(*20*) revealed broad association of RNAs with chromatin, it remains unclear how these associations relate to regulatory interactions between promoters and enhancers. Most existing RNA-chromatin proximity data is analysed at relatively coarse resolution, without directly connecting RNA binding to defined cis-regulatory loops. To bridge this gap, we integrated high-resolution enhancer-promoter maps obtained by promoter-capture Hi-C (HiCap) with RADICL-seq-based RNA-DNA association profiles (*20*). This approach allowed us to directly assess how RNA association at promoters and enhancers correlated with the formation, maintenance and rewiring of cis-regulatory interactions during neural differentiation, where human induced pluripotent stem cells (iPSCs) are differentiated into neural stem cells (NSCs) and then to neurons (NEU), described in detail in Lambolez et al. (in preparation). Here we demonstrate that promoter-associated RNAs are linked to the presence and stability of DNA–DNA contacts, whereas enhancer-associated RNAs modulate these contacts in relation to the RNA association state of the promoters – suggesting a coordinating role of RNA in cis-regulatory network dynamics.

## Methods

### Probe design

We downloaded RefSeq promoters (Dec 2, 2023, curated set) from the UCSC table browser(*21*). We only included genes with locus.type “gene with protein product”, “RNA, long non-coding”, “T cell receptor gene”, “unknown” and “immunoglobulin gene”. We removed olfactory receptor, ribosomal and histone protein genes (*22*). We designed probes for the *DpnII* fragment containing the transcription start sites. We designed separate probes for alternative promoters of a given gene if they were at least 1,200 bases apart from each other. We also designed probes for 2,003 variants associated with neurodevelopmental diseases (*23*). In total, we designed probes covering 5,940 Mb (115,770 probes) (purchased from Agilent Technologies with help from their design team).

SABER DNA FISH probes targeting promoters and corresponding HIRs were designed based on genomic coordinates extracted from HiCap datasets. For each coordinate, a 1.5-3 kb region centered on the locus was selected for probe set prediction using PaintSHOP (24). Probe sequences were further appended with distinct SABER primer sequences (25) to differentiate the promoter and HIR probe sets. All DNA oligos were ordered from IDT. SABER imager oligos conjugated to Alexa 488 or Alexa 647 were synthesized by Eurofins.

### Targeted chromosome conformation capture (HiCap) experiments

The Hi-C experiments on neural differentiation model were performed in duplicates. In brief, two millions of human induced pluripotent stem cells (iPSCs), neural stem cells (NSCs) and differentiated neurons (NEUs) per cell type and replicate were collected and crosslinked with 1% formaldehyde and subjected to Hi-C using the Arima HiC+ kit (Arima Genomics, cat. no. A510008) and the Arima Library Prep Module (Arima Genomics, cat. no. A303011). We hybridized Hi-C libraries to the probe set described above following Agilent SureSelect XT HS2 DNA System (G9983-90000) with slight modifications (*24*).

### SABER DNA FISH

*SABER DNA FISH probes were synthesized as described in* (*25*).

#### Primer-exchange reaction (PER)

We pre-mixed 1× PBS, 10 mM MgSO₄, 0.4 U/µL Bst LF polymerase (M0275S, NEB), 600 µM dATP/dCTP/dTTP (NTP-101, TOYOBO), 100 nM Clean G (Eurofins), and 1 µM SABER hairpin oligo (Eurofins), and incubated the reaction at 37 °C for 15 min. We then added probes containing the SABER primer sequence to the reaction and incubated the mixture at 37 °C for 120 min. After amplification, we heat-inactivated the reaction at 80 °C for 20 min. Finally, we purified the probe concatemers using a DNA Clean & Concentrator-25 kit (Zymo Research).

### DNA FISH

We cultured iPSCs and NSCs on round coverslips (18 × 18 mm) coated with iMatrix-511 (Matrixome), and we cultured NEUs on Poly-L-ornithine (Sigma-Aldrich)-coated coverslips. We washed the cells twice with 1× PBS and fixed them with 4% paraformaldehyde (Thermo Scientific) in 1× PBS for 10 min. After fixation, we washed the cells three times with 1× PBS and permeabilized them with 0.5% Triton X-100 (Sigma-Aldrich) in 1× PBS for 15 min. We then washed the cells twice and denatured them with 0.1 M HCl for 5 min.

Next, we rinsed the cells twice with 2× SSCT and treated them with 400 µg/mL RNase (Invitrogen) in 2× SSCT at 37 °C for 1 h. After rinsing twice with 2× SSCT, we incubated the cells in 2× SSCT containing 50% formamide for 5 min. We then transferred the coverslips to pre-heated 50% formamide in 2× SSCT in a 12-well plate and incubated them at 60 °C for at least 1 h.

For probe hybridization, we placed the coverslips cell-side down onto 10 µL hybridization buffer (67 nM PER PER concatemers, 50% formamide, 10% dextran sulfate in 2× SSCT) and sealed them with rubber cement (MP Biomedicals). We heated the slides at 80 °C for 3 min and then incubated them at 42 °C overnight.

The following day, we washed the cells three times in pre-warmed 2× SSC at 37 °C for 10 min each, and then we washed them in 0.1× SSC at 60 °C for 10 min. For imager oligo hybridization, we rinsed the cells with 1× PBS at 37 °C for 3 min and applied 1 µM fluorescent imager hybridization buffer (fluorescent imagers in 1× PBS + DAPI) directly to the coverslips. We incubated the cells in a humidified chamber at 37 °C for 1 h, washed them three times in pre-warmed 1× PBS at 37 °C, and mounted them in Antifade Vectashield.

### Microscopy and image analysis

We acquired confocal images on a Leica TCS SP8 confocal microscope using Leica Application Suite X (LAS X 3.5.7) and a 63× Plan-Apochromat NA 1.4 oil-immersion objective. We collected images using photomultiplier tubes (PMTs) and hybrid detectors (HyDs). The pixel size was 90–100 nm/px, and optical sections were acquired every 200 nm. For image analysis, we first segmented nuclei and identified FISH spots by applying median filter, Otzu thresholding, watershed modules in CellProfiler (v4.2.8). We then extracted the XYZ coordinates of FISH spot centroids and calculated the shortest promoter-HIR 3D distances in RStudio (v4.5.0).

### Sequencing and HiCap analysis

We sequenced HiCap libraries (2.5 nM pool) on Illumina NovaSeq X Plus (NovaSeqXSeries Control Software 1.2.0.28691) 10B mode flowcell with 300 cycles (2×150bp) configuration using one lane. We obtained 1603.46 clusters with 95.02% reads were above or equal to Q30. We mapped the reads using BWA-mem2 (v2.2.1) (*25*), processed the pairs and removed the duplicate reads using Pairtools (v1.1.3) (*26*). We then used HiCapTools (version 1.3.2) (*27*) to call for significant genomic interactions for each replicate. We required minimum five supporting pairs and a minimum Bonferroni-corrected p-value of 0.05 to be counted as a significant interaction. We removed interacting regions overlapping blacklisted regions (downloaded from https://hgdownload.soe.ucsc.edu/gbdb/hg38/problematic/encBlacklist.bb) (*28*).

### RADICL-seq data processing

RADICL-seq data produced from the neural differentiated set is described in detail in Lambolez et al (in preparation). We took significant RADICL-seq interactions (FDR < 0.01) and removed regions overlapping blacklisted regions (downloaded from https://hgdownload.soe.ucsc.edu/gbdb/hg38/problematic/encBlacklist.bb) (*28*). We also removed interactions where the RNA is interacting with its host locus (i.e. self-interactions). We merged overlapping DNA regions but did not perform any binning on the dataset. We also divided the RADICL-seq dataset based on the source of the RNA reads as intronic or exonic reads or with no respect to the source of the RNA (All). We additionally classified RADICL-seq dataset based on the distance RNA-DNA interactions spanned: cis interactions are counted as those where the genomic locus of the source RNA and the target DNA is less than 1.25 Mb away, trans interactions were those where the source RNA and target DNA were on different chromosomes or the distance spanned longer than 1.25 Mb if they were on the same chromosome. The average width of target DNA is shown in Supplementary Figure 1a. We assigned the number of RNA reads originating from a given RNA to a given target DNA as the weight of the interaction, the distribution of weights is shown in Supplementary Figure 1b.

### HiCap-RADICL-seq network generation

We processed each replicate for each cell type separately. First, we overlapped the DNA part of the RADICL-seq data to HiCap interactors: promoters and promoter-interacting regions (PIRs). We then removed the rest of the RADICL-seq data which did not involve any HiCap regions but kept all the HiCap interactions. This generated interactions between the HiCap regions and RNA transcripts (RNA-DNA) as well as the interactions between the HiCap regions (DNA-DNA). We generated interaction lists by combining each of the two replicates of HiCap with each of the two replicates of the RADICL-seq data to avoid spurious contacts due to merging between the replicates (HiCap replicate 1 / RADICL-seq replicate 1, HiCap replicate 1 / RADICL-seq replicate 2, HiCap replicate 2 / RADICL-seq replicate 1, HiCap replicate 2 / RADICL-seq replicate 2) in each cell type, totaling to 12 interaction sets (4 for each cell type). The interaction statistics were calculated over the four replicates. For network dynamics and individual RNA interaction analyses, we only took the interactions present in all four replicates. We converted each interaction dataset to a network using the igraph package (v 2.2.1) (*29*) available via R. The degree (number of interactions) and clustering coefficient (the ratio of the number of connected neighbors of a node to all possible connections between the neighbors) is calculated by the igraph package.

### Network Randomisation

To determine the probability of contacting the same RNA for the two interaction regions in HiCap data, we generated randomised counterparts of each of the 12 networks. We did not randomise the HiCap interactions and we processed each chromosome separately. We randomised the interacting RNA of a HiCap interactor while respecting the chromosome and number of connections of each node. We used Networkit (v 11.0) (*30*) package and used the Global curveball randomisation scheme, which considers neighborhoods of the randomly drawn node pairs (*31*) while preserving the degree of each node.

### RNA interaction effect on DNA-DNA interactome dynamics

To assess the impact of RNA association on DNA–DNA interaction dynamics, we constructed networks by integrating all the RADICL-seq data (independent of their overlap with HiCap interacting regions) with the HiCap data. Using the RADICL-seq data, we then examined the RNA association status of every HiCap region, regardless of whether it participated in a DNA-DNA interaction across all tested cell types. This allowed us to determine the RNA association status of promoters and PIRs in each cell type, even in the absence of a DNA-DNA interaction. For all analyses, only replicated RADICL-seq and HiCap interactions were considered. DNA-DNA interaction dynamics were classified as lost, gained, or stable based on their reproducibility across both HiCap replicates.

### Normalization of Gain and Loss Interaction Enrichment Relative to Stable Interaction Enrichment

To quantify how promoter and PIR RNA–association dynamics influence changes in DNA–DNA interaction status, we calculated enrichment scores for gained, lost, and stable interactions for each promoter–PIR RNA-change category. Each RNA-change category is defined by a pairwise transition in RNA-association state between cell types (PromoterRNAchange × PIRRNAchange), resulting in four possible states for each side: Gain, Loss, Stable(+), and Stable(−).

#### Step 1: Enrichment of interaction outcomes relative to a baseline

For every RNA-change category, we computed the proportion of gained, lost, or stable DNA–DNA interactions:

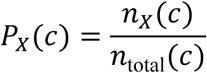

where

- 𝑋 ∈ {Gain,Loss,Stable},
- 𝑐 is an RNA-change category,
- 𝑛*_x_*(𝑐) is the number of interactions with outcome 𝑋, and
- 𝑛_total_(𝑐) is the total number of interactions in that category.

Enrichment values were then computed relative to a biologically neutral baseline category:

**Stable(−)_Stable(−)**

(no RNA association at either promoter or PIR in either cell type). Baseline enrichment for outcome 𝑋 was:

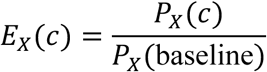

Enrichment values were set to 1 for the baseline category. For each RNA-change category and interaction outcome, enrichment significance was evaluated relative to the baseline category using Fisher’s exact test on a 2×2 contingency table:

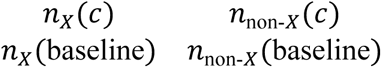

False discovery rate was controlled using Benjamini–Hochberg correction. Significance (FDR-adjusted) was represented on plots as ∗ (FDR < 0.05), ∗∗ (FDR < 0.01), or ∗∗∗ (FDR < 0.001).

### Quantification of interaction dynamics using Jaccard Index

We took only the DNA-DNA and DNA-RNA interactions present in all four replicates in each cell type. We calculated the Jaccard similarity index for each node (a) by comparing their connections in each cell type using the following formula:

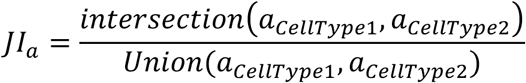

where intersection refers to the neighbors that stayed the same between Cell Type 1 and Cell Type 2, and union refers to the total number of unique neighbours of node a in Cell Type 1 and Cell Type 2. We performed these operations for each node by contrasting NSC vs iPSC, NEU vs NSC and NEU vs iPSC.

### Determining the regulatory targets of RNA

We used the igraph package (v 2.2.1) (*29*) available in R to first generate ego networks of each node (only the connections of the node in concern) with order = 2, to take both direct interacting partners and the interacting partners of its interactors for each RNA. The target genes of RNAs associated with promoters were defined as the genes to which the corresponding promoters belong. The target genes of RNAs associated with PIRs were assigned based on HiCap interactions of the corresponding PIRs. If RNA was interacting to both promoter and PIR of the gene, then it is classified as Prom/PIR interactor.

For the UMAP clustering, we calculated the Jaccard similarity index to quantify overlap of target genes of each RNA using igraph’s “similarity” function (method = “jaccard”, mode= “all”). We only included RNAs bound to at least 10 promoters and/or PIRs in this analysis.

### Community Detection

We used community detection to group nodes that are more connected to each other than to the rest of the network into communities or modules. This method coarse-grains large interaction maps into compact, interpretable units and helps relate network structure to biological function (e.g., groups of promoters, regulatory elements, and RNAs that tend to act together). In practical terms, communities capture regions in the network of dense connectivity and reduce noise by focusing analyses on coherent subgraphs rather than on individual nodes and links. There exists a vast library of computational tools for finding such groups. We used Infomap (version 2.8.0), an information-theoretic method that partitions the graph by minimizing the so-called map equation (Infomap effectively groups nodes that a fictious random walker revisits many times together (*32*)). We chose Infomap because it has been thoroughly tested and performs well on biological networks, for example associated with gene regulation (*33*, *34*). We ran Infomap (default settings) on our weighted graphs with multiple restarts to find the best partition.

## Results

We performed targeted chromosome conformation capture (HiCap) on a neural differentiation series derived from induced pluripotent stem cells (iPSCs), sampled at three stages: iPSCs, neural stem cells (NSCs), and neurons (NEUs), each with two biological replicates (*35*). The HiCap probe set targeted 26,394 promoters corresponding to 22,505 genes, 2,003 neurodevelopmental disease–associated SNVs, and 5,118 negative control regions (Supplementary Table 1). Across the six HiCap libraries, we generated 545.3 million unique read pairs, of which 71% mapped in cis; on average, 45% (221.7 million) spanned distances greater than 40 kb (Supplementary Table 2). For each sample, we constructed a sample-specific background interaction frequency distribution using negative control regions and identified statistically significant interactions by comparing observed frequencies against the respective sample-specific background (*27*). This analysis yielded 334,990 significant interactions (Bonferroni-adjusted p < 0.05 and supported by ≥5 read pairs; Supplementary Table 3) (Figure 1a). The average interaction distance was 174.4 kb (Figure 1b), and replicates of the same cell type showed highly similar interaction counts (Figure 1c, and Supplementary Figure 2a–c).

**Figure 1.**
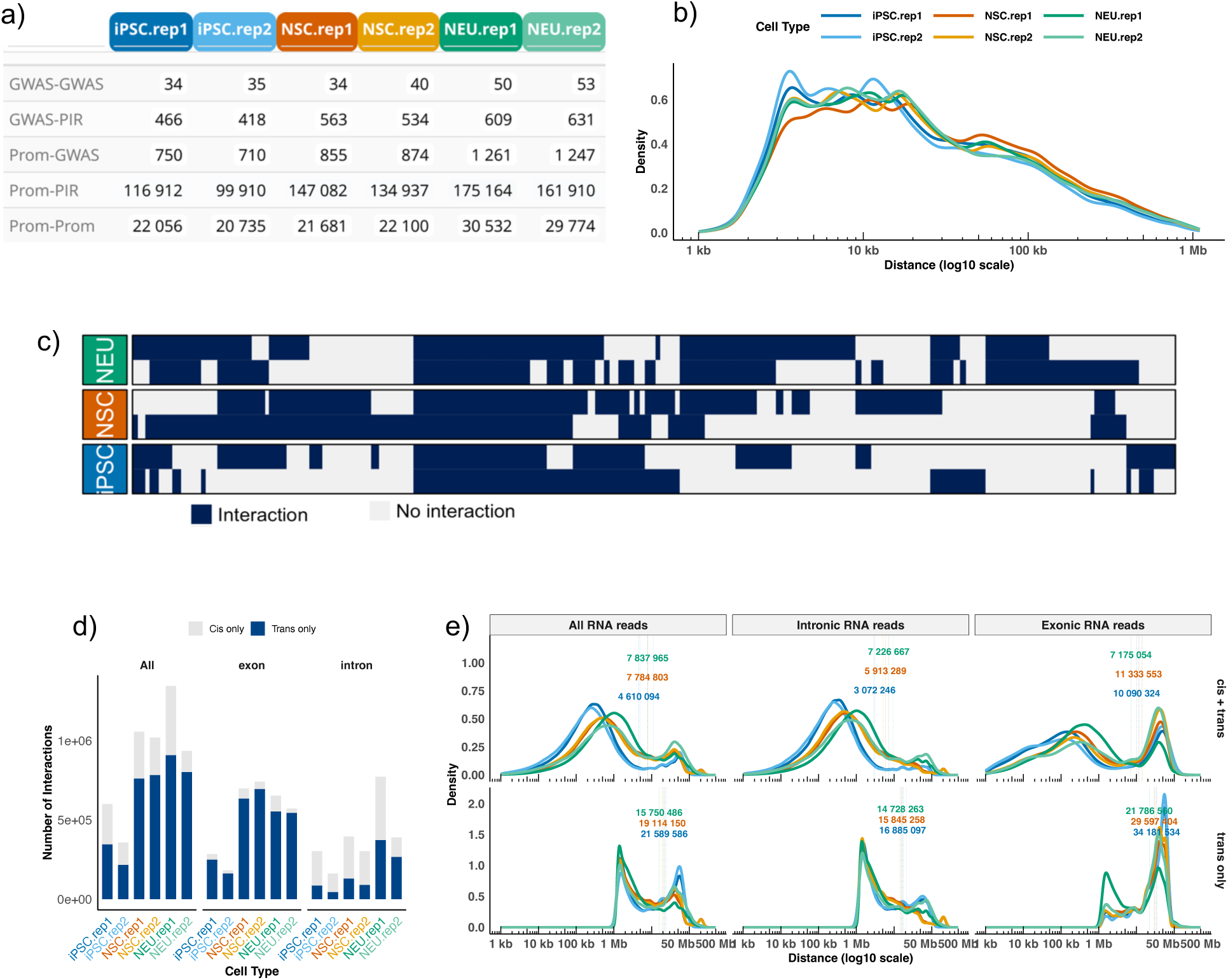
**a)** The number of interactions in each HiCap dataset, GWAS: GWAS variants associated with neurodevelopmental diseases targeted in the HiCap probe set, Prom:Promoter, PIR: promoter interacting region; **b)** The distance distribution of HiCap interactions in different cell types and replicates; **c)** The binary heatmap of each HiCap sample (grouped by cell type), only interactions in chr21 is shown for illustration, blue denotes the presence of an interaction; **d)** The number of cis (grey) and trans (blue) interactions in each RADICL-seq dataset, separated either using all RNA reads, only exonic or intronic RNA reads; **e)** The distance distribution of the genomic locations of target and source regions in RADICL-seq data, separated by the source RNA type (all, exonic or intronic). The top panel shows all associations, the bottom panel shows the trans-associations where the target and source genomic loci are on the same chromosome.

We next analyzed RNA–DNA proximities using RADICL-seq (RNA And DNA Interacting Complexes Ligated and sequenced) (*20*) data generated in the same cell types (Lambolez et al., in preparation). RADICL-seq identifies DNA regions (RD-DNAs) that are in proximity to RNA transcripts. To identify statistically significant RNA–DNA proximities, hereafter referred to as associations to distinguish them from statistically significant DNA-DNA interactions, we applied the CHICANE method (*35*) (FDR < 0.01). We retained only significant RNA–DNA associations and excluded those between each RNA and its own genomic locus, which we considered “self-interactions.” This resulted in a total of 1,910,566 RNA–DNA associations (Supplementary Table 4, Supplementary Figure 2d). We further categorized these into six sets based on RNA origin (any, intronic, or exonic) and the genomic distance between the RNA locus and its DNA target. Trans associations were defined as those occurring either between different chromosomes or between loci separated by at least 1.25 Mb (see Methods). Trans associations predominated across all cell types (Figure 1d). Exonic RNA–DNA associations were predominantly trans and spanned greater distances largely due to extensive long-range contacts made by the mono-exonic MALAT-1 gene (Figure 1e).

### RNA-associated regions are enriched for gene regulatory interactions

To better understand the functional roles of caRNAs detected by RADICL-seq, we assessed how many RNA-interacting regions overlap individual gene regulatory elements such as promoters and promoter-interacting regions (PIRs) by combining our HiCap and RADICL-seq datasets. To move beyond pairwise interactions, we constructed integrated networks incorporating both DNA-DNA interactions and RNA–DNA associations (see Methods). For each cell type, we overlapped HiCap interacting regions (HIRs) with RNA-associated DNA regions identified by RADICL-seq (hereafter RD-DNAs).

We generated four networks per cell type by combining two HiCap and two RADICL-seq replicates, resulting in a total of 12 networks (see Methods). This integration yielded 2,186,538 combined interactions involving 280,790 genomic regions and 11,253 RNA transcripts (Supplementary Table 5). Interaction numbers were consistent across replicates (Supplementary Figure 2e), with 722,345 interactions shared by all replicates of a given cell type (Supplementary Figure 2f, Supplementary Table 6). Promoters with HiCap interactions were associated with higher numbers of RNAs and this pattern persisted even when restricting to trans RNA–DNA associations (Figure 2a, Supplementary Figure 3a). In contrast, RD-DNA regions were associated with higher number of RNAs compared to PIRs on average (Figure 2b, Supplementary Figure 3b), confirming the multi-faceted role of chromatin-associated RNAs beyond canonical transcriptional regulation(*12*, *36*, *37*).

**Figure 2.**
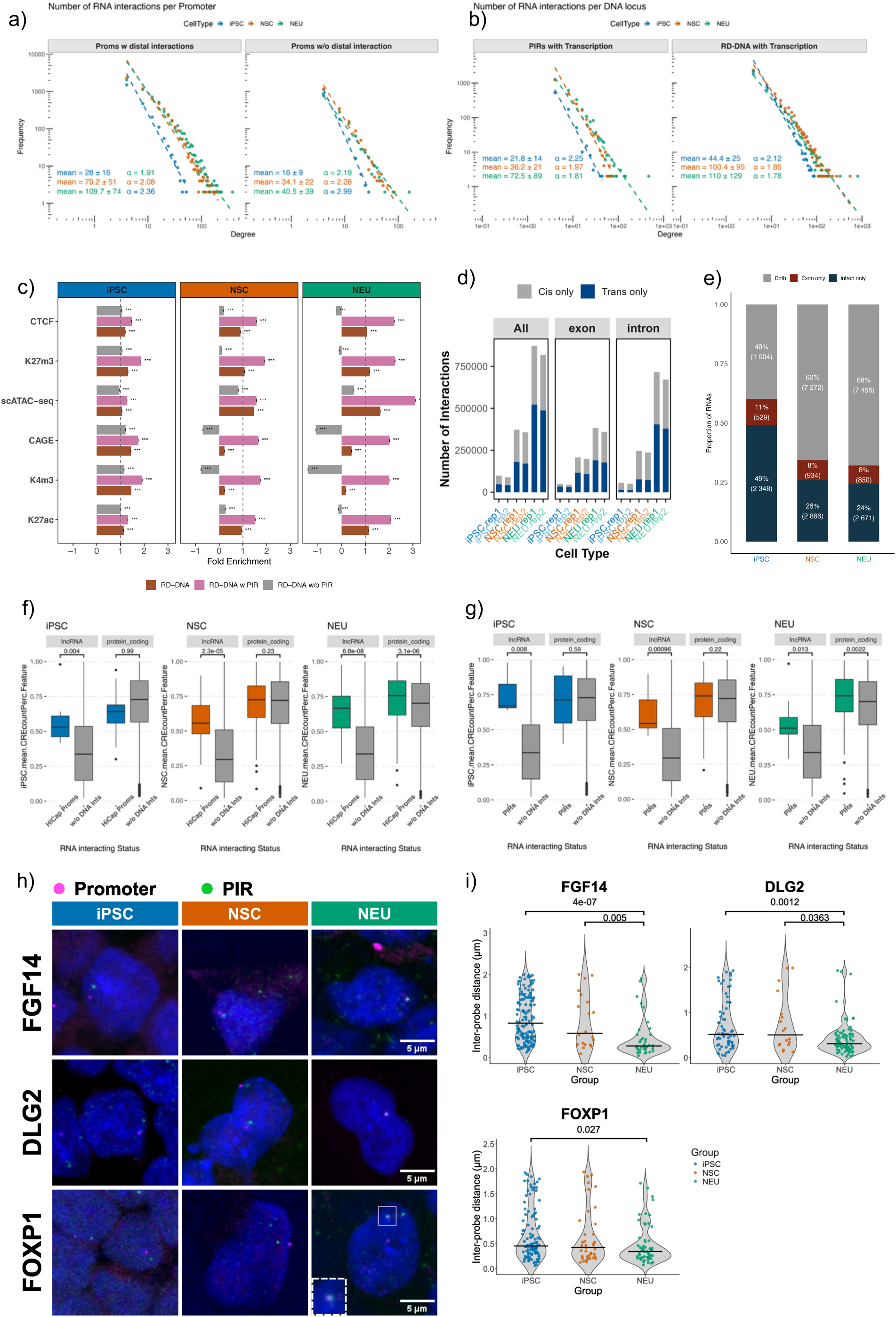
**a)** Degree distribution of HiCap promoters and promoters with no distal interaction, x-axis denotes the degree (the number of interacting RNA transcripts) of promoters, y-axis corresponds to the number of promoters with that degree. The slope of the line fits are shown on the top right; **b)** The degree distribution of HiCap interacting regions (PIRs, excluding promoters) and DNA regions (excluding all promoters and PIRs) containing CAGE peaks; **c)** The fold enrichment of HiCap regions in terms of their RNA interaction status for H3K27Ac, H3K4me3, CAGE, single-cell ATAC-seq, H3K4me3 and CTCF binding sites; **d)** the number of RNA-DNA interactions of which DNA part overlaps with a HiCap region stratified by RNA source and RNA-DNA distance span; **e)** the percentage of RNAs engaged interactions via exonic, intronic or both segments; **f)** The expression percentile of lncRNA or protein coding RNA transcripts interacting with HiCap promoters or promoters with no distal interaction, only trans RNA-DNA interactions is used; **g)** The expression percentile of lncRNA or protein coding RNA transcripts interacting with PIRs only non-promoter DNA.

We observed a 9.3-fold enrichment (p = 0) for HIRs within RNA-interacting DNA regions (6.17%, 25.3 Mb). As expected (*16*), both RD-DNA regions and HIRs were enriched for regulatory elements, with regions overlapping both datasets showing stronger enrichment for activating marks and open chromatin, particularly in NSCs and NEUs (Figure 2c).

RD-DNA regions overlapping with HIRs were more likely to associate with an RNA in *cis* (Figure 1d and 2d) and the source of RNA for a substantial number of DNA-associated transcripts was derived from both exonic and intronic reads (Figure 2e). Finally, RNAs interacting with HIRs were more highly expressed than those interacting with non-HIR regions, irrespective of gene type (Figure 2f). When restricting to only *trans* RNA–DNA interactions, only lncRNAs interacting with HIRs showed significantly higher expression, except in NEUs where both lncRNAs and protein-coding RNAs were more highly expressed (Figure 2g). To confirm this, we performed SABER DNA FISH to validate HIR formation across iPSC-to-NEU differentiation. We selected three HIR pairs (promoters of FGF14, DLG2, and FOXP1 and their corresponding interacting loci) that were identified as interacting with intronic RNAs specifically in NEUs. By measuring the distances between promoter signals and their corresponding PIR signals, we found that distances for the FGF14, DLG2, and FOXP1 HIR pairs were reduced by 550 nm, 205 nm, and 110 nm, respectively, in NEUs compared with iPSCs. In addition, all HIR pairs exhibited shorter distances in NEUs than in NSCs, although the FOXP1 pair did not show a statistically significant difference. Distance distribution analyses also revealed a higher proportion of cells with shorter HIR distances in NEUs, indicating an increased frequency of HIR contacts. Together, these results suggest that intronic RNA-mediated HIR formation is a cell-type-specific event during neuronal differentiation (Figure 2h and 2i).

Next, we classified each promoter (Prom), GWAS variant (GWAS), and promoter-interacting region (PIR) according to whether or not they are associated with an RNA as informed by RADICL-seq, using the following nomenclature: Prom|RNA(−) vs. Prom|RNA(+), GWAS|RNA(−) vs. GWAS|RNA(+), and PIR|RNA(−) vs. PIR|RNA(+) for each cell type and replicate (Figure 3a). PIR|RNA(+) regions were supported by a significantly higher number of HiCap read pairs compared to PIR|RNA(−) regions (1.18-fold increase; p < 8.2 × 10⁻¹⁰, Welch two-sample t-test). This difference was even more pronounced when the interacting promoter itself was RNA-associated (1.23-fold; p < 2.2 × 10⁻^16^, Welch two-sample t-test). RNA-associated HIRs [HIR|RNA(+)] engaged in a higher number of DNA–DNA interactions compared to HIR|RNA(−) (Figure 3b; Supplementary Fig. 3c). Interacting DNA regions were more likely to connect to the same RNA transcript(s), as indicated by a higher clustering coefficient, which measures connectivity between the neighbors of each node (Figure 3c; Supplementary Fig. 3d see Supplementary Material). This was also confirmed by comparing the chance of observing the same pattern in corresponding randomized networks where the degree and HiCap interactions were kept the same and the RNA-DNA associations were randomized per chromosome (see Methods) (Figure 3d). These findings support recent observations that caRNAs can act as scaffolds for organizing and stabilizing DNA-DNA contacts(*36*, *38*, *39*).

**Figure 3.**
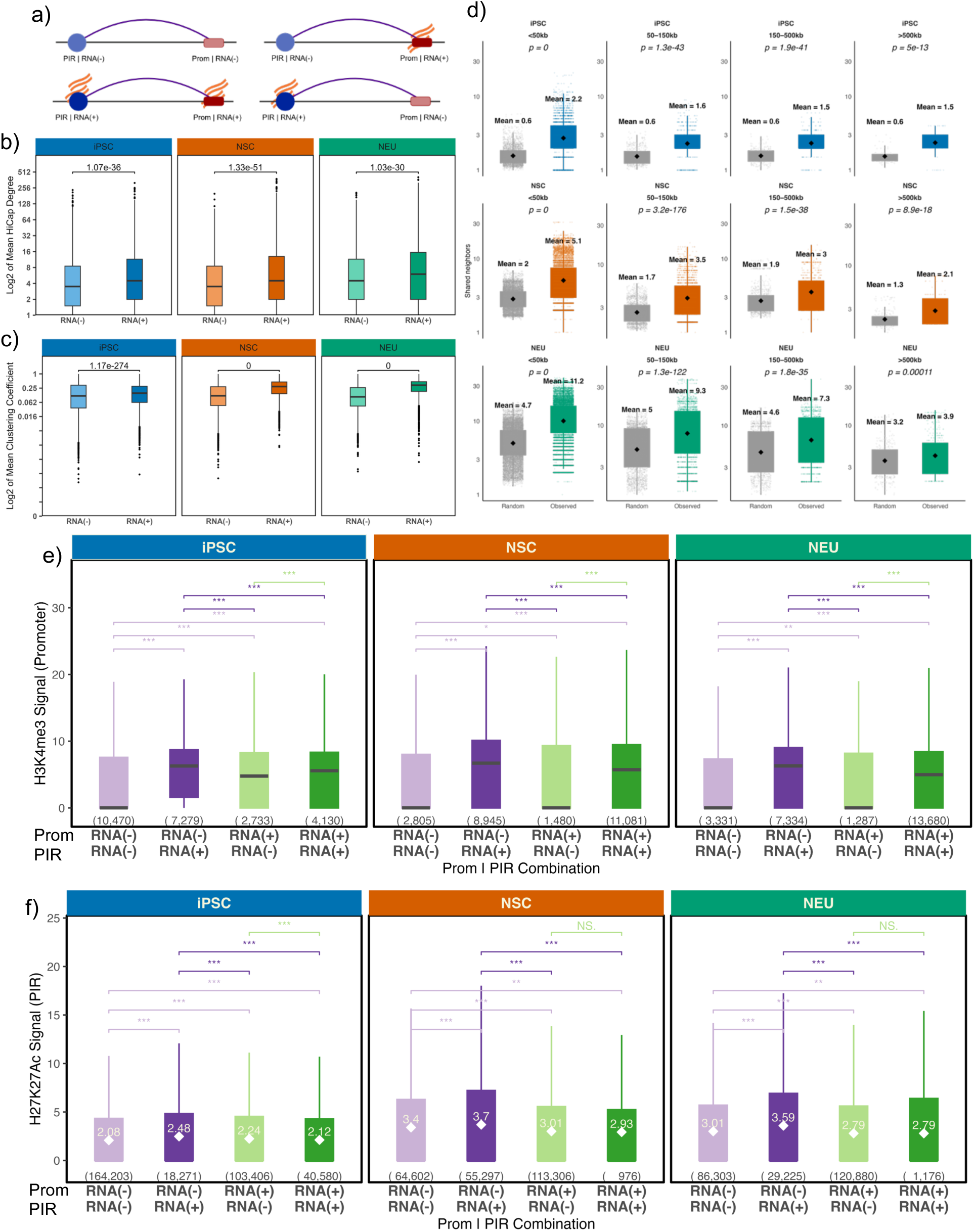
**a)** A sketch of interacting (purple arcs) promoter (red) and PIR regions (blue) with their RNA (orange curved lines) interaction status combinations; **b)** the number of PIR connections; **c)** the clustering coefficient of promoters with respect to their RNA interaction status; **d)** the average number of shared RNAs per HiCap interaction in the observed and random networks stratified by cell type and the genomic distance spanned by the HiCap interaction; **e)** The H3K4m3 levels of promoters; **f)** H3K27Ac levels of PIRs in different combinations: gray: both promoter and PIR RNA(−), yellow: only PIR is RNA(+), purple: only promoter is RNA(+), both promoter and PIR is RNA(+).

Promoters interacting with PIR|RNA(+) regions had higher active chromatin mark levels than those interacting with PIR|RNA(−) regions, and Prom|RNA(−) interacting with PIR|RNA(+) had higher levels of active chromatin marks than Prom|RNA(+) interacting with PIR|RNA(−) (Figure 3e). Furthermore, PIR|RNA(+) regions showed higher H3K27ac levels compared to PIR|RNA(−) regions (Figure 3f). Overall, these results confirm that RNA association status of enhancers may in part modulate their own chromatin activity as well as that of their interacting promoters (*17*).

### RNA association dynamics of regulatory elements coincide with DNA-DNA interaction changes across cell types

We next investigated whether RNA association of HIRs coincided with changes in their DNA-DNA interaction dynamics during differentiation. We compiled RNA association and DNA interaction status of all HIRs, retaining only interactions detected in all replicates and occurring in *trans* to prioritise changes more likely driven by functional RNA binding rather than local RNA diffusion (see Methods). We then identified HIRs that gained or lost RNA associations between consecutive cell types (iPSC vs. NSC and NSC vs. NEU), as well as those whose RNA association status remained unchanged (see Methods). For each category, we compared the fraction of stable, lost, and gained DNA-DNA interactions across the consecutive cell types. Changes in RNA association were strongly related to DNA-DNA interaction outcomes (Figure 4a-c) (p = 0, Chi-square test). Cases in which both promoter and PIR gained RNA association were markedly enriched for gained interactions (1.6-fold, padj = 0), similarly loss of RNA association on both promoter and PIR was enriched for interaction loss between them (1.5-fold, padj = 8e-12) (Figure 4d-f). Stable DNA-DNA interactions were most enriched when both promoter and PIR maintained their RNA association state across cell types (36,650 interactions, 1.9-fold, padj = 1.8e-42) (Figure 4e). The strongest enrichment for DNA-DNA interaction gain occurred when PIRs gained RNA association while promoters remained RNA-associated in both cell types (43,085 interactions, 1.95-fold, padj = 0). RNA association gain on promoters while PIR stayed associated with RNA were also strongly correlated, though this category was smaller (5,585 interactions, 1.85-fold, padj = 8.3e-153). The loss of RNA association at PIRs were consistently enriched for the loss of DNA interaction with promoters independent of promoter RNA association status (Figure 4f). However, the loss of promoter RNA association was more likely to coincide with interaction gain when the PIRs gained RNA association (418 interactions, 1.5-fold, padj = 1.1e-19). These findings indicate that RNA association dynamics, particularly at PIRs closely coincide with and may help the coordination of DNA-DNA interaction changes, rather than solely reflecting recruitment to pre-existing chromatin loops.

**Figure 4.**
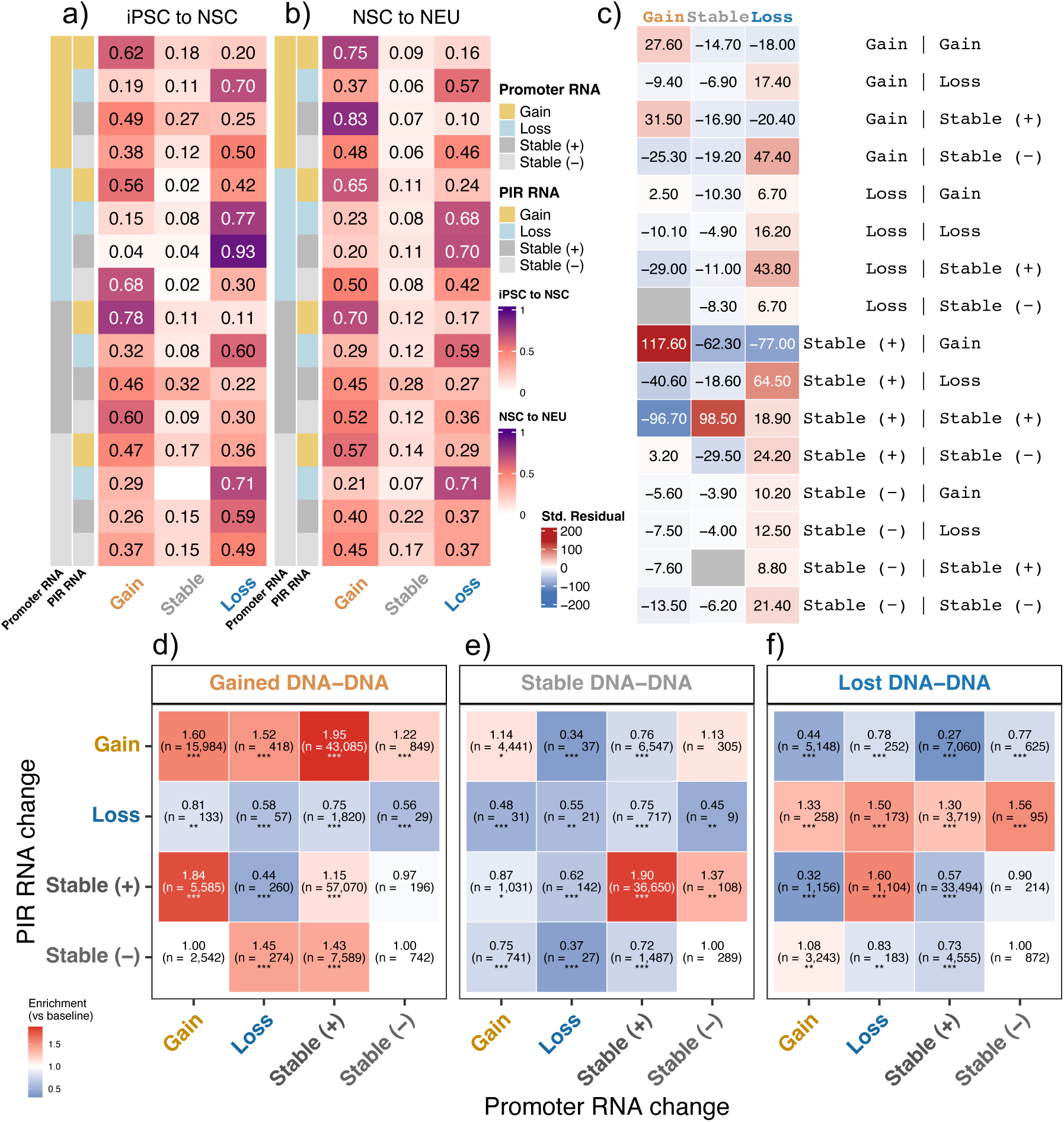
The fraction of gained, lost and stable HiCap interactions in during; **a)** iPSC to NSC; **b)** NSC to NEU transition with respect to the RNA interaction profile of the HIRs; Gain: gained, Loss: lost an RNA interaction in the consecutive cell type; Stable (−): no RNA interactions in either cell types, Stable(+): the same RNA interactions in both cell types; **c)** The heatmap showing standardized Pearson residuals from the chi-square test, red: observed>expected, blue: observed<expected. The fold-enrichment of **d)** gained or **e)** stable **f)** lost interactions with respect to RNA association dynamics of promoters and PIRs. The fold ratios are calculated by normalizing with respect to the number of regions with no RNA associations (see Methods).

### RNA associations with DNA cis-interactome are highly dynamic during differentiation

We next examined the dynamics of RNA associations involving promoters and PIRs, again considering only those interactions detected in all four replicates. Differentiation was accompanied by a pronounced increase in RNA interactions: only 10,023 features and 6,753 edges were shared across all cell types, pointing to the highly dynamic nature of the RNA–chromatin interactome (Figure 5a–b). Specifically, 757, 1,818, and 2,056 RNA transcripts associated with HIRs exclusively in iPSC, NSC, and NEU cells, respectively (Figure 5c). Of the 18,346 HiCap promoters, 1,170 (6.3%) did not have any RNA associations in any cell type, while 3,702 (28%) were RNA-associated in all cell types. Strikingly, 13,579 (74%) promoters displayed cell type–specific RNA association status.

**Figure 5.**
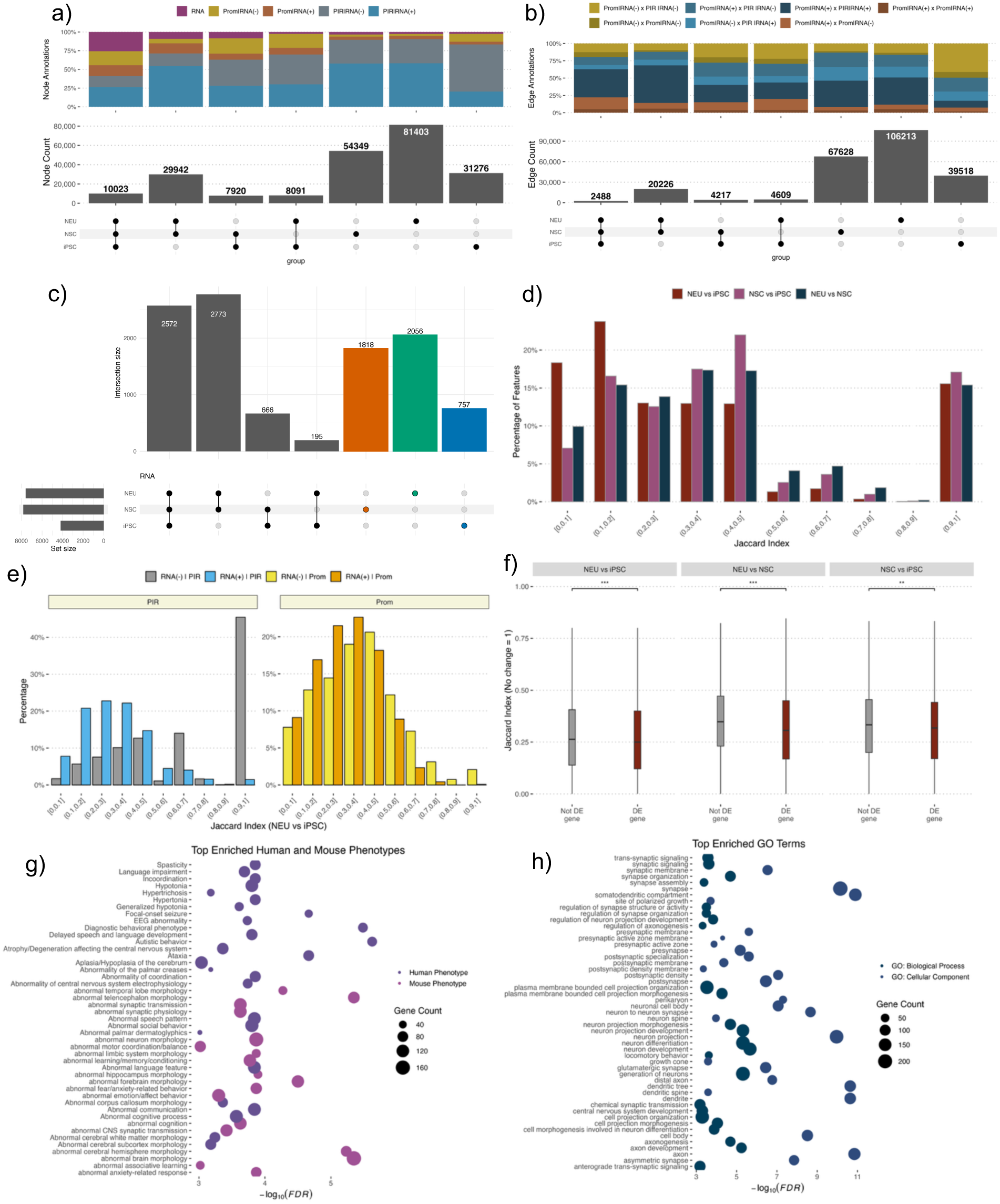
The number of **a)** feature (node) **b)** interaction (edge) **c)** RNA transcript distribution across different cell type combinations. The distribution of jaccard indexes **d)** across cell type comparisons; **e)** relative to the RNA interaction status of promoter-interacting regions (PIRs) or promoters; **f)** of the differentially vs non-differentially expressed genes; The list of enriched **g)** human and mouse phenotypes **h)** GO terms for the DE genes with a jaccard index < 0.1 in iPSC vs NEU cell types. All DE genes are used as the background set while performing enrichments.

Given this extensive cell type specificity, we next asked whether promoters that altered their RNA association status across differentiation belong to genes relevant for neural lineage progression. To address this, we compared promoter-associated RNA sets across cell types and quantified their similarity using the Jaccard index (*40*). For each promoter, we computed the Jaccard Index between its associated RNA sets across cell types, providing a normalized measure of overlap that account for both the number and identity of associated RNA. To focus on associations more likely to reflect functional RNA binding rather than transcriptional proximity due to local RNA diffusion, the analysis was restricted to only *trans* RNA–DNA associations (see Methods). The Jaccard index ranges from 0 indicating complete rewiring of RNA association, to 1, indicating no change between conditions (Supplementary Material) (*41*). Most features exhibited substantial RNA association changes during differentiation (Figure 5d). Promoters and PIRs that engaged in RNA associations were significantly more dynamic (i.e. more likely to be rewired) than their RNA-negative counterparts (Supplementary Figure 4a). Even among features that maintained the same RNA association status between cell types, those that were RNA-positive in both cell types displayed greater rewiring of their DNA interaction partners compared to RNA-negative elements (Figure 5e). Promoters of differentially expressed genes also showed significantly higher interaction rewiring (Figure 5f). Finally, comparative gene enrichment analysis, using all differentially expressed genes as the background, revealed that differentially expressed genes with the strongest interaction changes (Jaccard index < 0.1; 913 genes) were significantly enriched for neural differentiation–related functions, whereas differentially expressed genes with stable interactions (Jaccard index > 0.4; 1,340 genes) showed no enrichment (Figure 5g–h).

### RNA transcripts associate with enhancers to amplify the up or down-regulation of target genes

We showed that *DNA-DNA* interaction dynamics are closely associated with changes in RNA association during neural differentiation. We next asked whether the identity of the RNA transcripts associated with HIRs influences transcriptional output. For this analysis, we again considered only replicated interactions in all four networks of any cell type retaining only *trans* RNA-DNA interactions. As expected, most associated RNAs were cell-specific and enriched for being differentially expressed (Figure 6a). To characterize RNA association patterns, we counted for each RNA the number of non-HIR regions (RD-DNA only), PIRs and promoters it associated in each cell type. We also defined a separate category for RNAs that associated both the promoter and the PIR of the same gene (“Promoter & PIR”). As differentiation progressed, RNAs became significantly more likely to contact with PIRs and less likely to interact with non-PIR regions (Figure 6b, Supplementary figure 4b).

**Figure 6.**
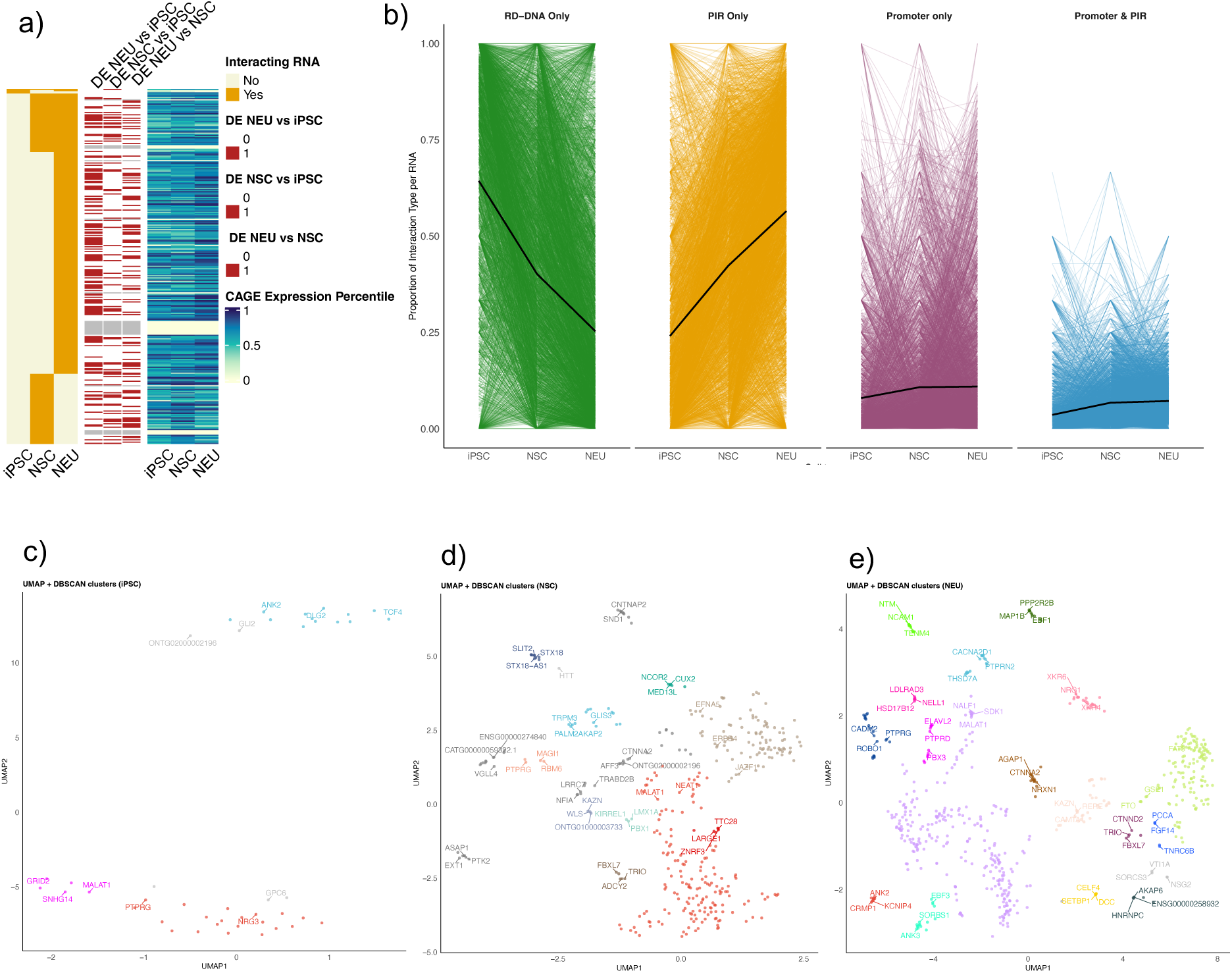
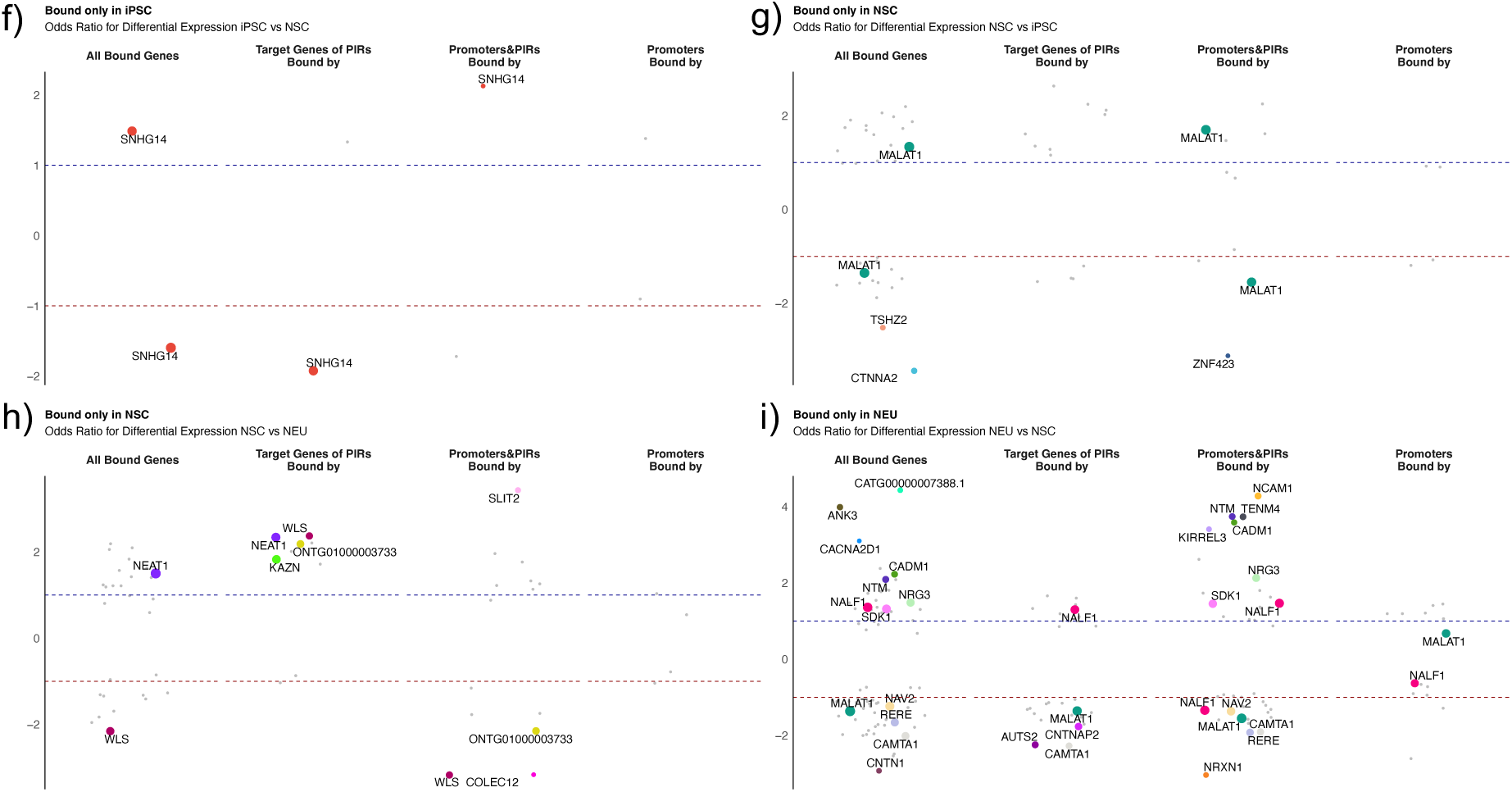
**a)** the interaction, differential and CAGE expression profile of 309 RNAs (4, 113 and 246 transcripts in iPSC, NSC and NEU, respectively) which only interacted via exonic reads and in trans manner (due to figure space limitations), Of these, 63% (139) were differentially expressed in at least one time point, however, there were many RNAs with similar RNA expression levels across cell types with different DNA binding pattern; **b)** spaghetti plot of the proportion of changes of each RNA, each line traces the change in the proportion of PIR-only, RD-DNA only, promoter only or promoter/PIR (of the same gene) of a single RNA across the cell types; **c-e)** UMAP clustering of interacting RNAs in each cell type based on the commonality of the promoters/PIRs they bind calculated by Jaccard similarity index, only those bound to at least 10 regions are taken into account for clustering, colors indicate separate clusters and only RNA with high degree are labeled; **f-i)** The odds-ratio whether genes bound by the RNA of interest in a cell type specific manner are enriched among differentially upregulated (positive) or downregulated (negative) genes. Only those RNA with odds ratio > 1.3 and FDR < 0.1 is labelled. Promoters/PIRs corresponds to genes whose promoters and PIRs interacting with that promoter bound to the RNA(s). Only consecutive cell types (iPSC vs NSC and NSC vs NEU) are considered.

We then compared the RNAs associated with each promoter in each cell type to determine whether promoters share similar RNA-binding profiles. For each cell type, we computed the Jaccard similarity between every RNA-RNA pair based on their DNA targets to find RNAs associated to similar set of DNA regions (see Methods) and clustered RNAs based on their DNA target similarities. This analysis revealed distinct groups of RNAs with shared DNA association profiles in each cell type (Figure 6c-e). For example, *SNHG14* (Small Nucleolar RNA Host Gene 14) is a lincRNA located in Prader-Willi and Angelman syndrome imprinting locus (*42*) and have a neuron-specific imprinting pattern. Its antisense transcript is only expressed in neurons from the paternal allele (*43*). In our dataset, SNHG14 was bound 226 and 1,159 HIRs in iPSC and neurons but did not contact any region in NSCs. In iPSCs, it was bound to either promoters or PIRs of 114 genes, in neurons, this number was 63 and there was no overlap between its target genes in iPSC and neurons. The genes bound by SNHG14 in iPSCs were slightly enriched for transcription factors (21 genes, FDR = 3.39E-3) but no functional enrichment was present for the genes bound by SNHG14 in neurons.

Finally, we assessed whether promoters or PIR-target genes associated with a given RNA or RNA set were enriched for differential expression. We calculated the Jaccard similarity index for each RNA to compare its DNA-association pattern between consecutive cell types: i.e., iPSC vs NSC and NSC vs NEU cells (see Methods) to identify RNAs with substantial rewiring (Jaccard Index < 0.5). For each such RNA, we then assessed whether the promoters or the promoters targeted via PIRs were more likely to be differentially expressed. Indeed, we identified a strong effect for 29 RNAs (43.9%, out of 66 dynamic RNAs associated with at least 30 promoters/PIRs in total). Interestingly, there was no enrichment for differential expression when an RNA exclusively interacted with promoters. However, for many RNAs, depending on the RNA and the cellular context, genes they were associated either directly via promoters or indirectly via PIRs were more likely to be differentially expressed (FDR < 0.1) (Figure 6e-h). Target genes of RNAs such as KAZN, NEAT1 or NRG3 were more likely to be upregulated, whereas the target genes of RNAs such as AUTS2 or NAV2 were more likely to be downregulated. For other RNAs, such as SNHG14, MALAT1 or NALF1, the target genes were more likely to be differentially expressed but in no specific direction.

### RNA increases nonrandom connectivity across neural differentiation

In our interaction graphs, nodes represent genomic elements (promoters, PIRs, and RNA-associated DNA targets (RD-DNA)) linked by DNA–DNA or RNA–DNA contacts. To capture higher-order topologies, we sought to identify *communities* that are groups of nodes that interact more with each other than with the rest of the network. We extracted these communities using Infomap (see Methods) (*44*). In the iPSC, NSC, and NEU networks, again using only the *trans* and replicated data, we detected 7,279, 7,255, and 6,589 communities, respectively. Of these, 2.8%, 4.6%, and 6.3% are large communities, defined as having ≥100 nodes. The rest are small with less than 100 nodes. Because contact patterns of nodes within each community might change during differentiation, a node may remain in the same community, move to a different one or switch category (large vs small). It may also be absent when no qualifying links are detected in each cell type, despite appearing in the others (Figure 7a). To follow this regulatory architecture across differentiation, we tracked each node’s community assignment across cell types (Supplementary Figure 5). The community size and the number of communities increased upon differentiation. The fraction of nodes belonging to large communities increased from 18.5% in iPSC to 27.2% in NSC and 35.2% in NEU and the number of large communities increased from 202 to 331 to 418, respectively (Figure 7b). Importantly, the increase in the number of large communities was not simply a consequence of the increased number of RNA associated nodes in later stages as after normalising the number of connections by total RNA node counts, the proportion of nodes joining large communities still grew disproportionately (Supplementary Figure 6). Moreover, the driver of the increase in both the size and number of large communities was not the frequent switching of nodes between existing communities (which remains low at ∼7%) but rather the appearance of new nodes and the merging of existing DNA-DNA communities mediated by RNA connections. Newly appearing DNA nodes often attached via RNA-linked contacts, bringing clusters of nodes into existing communities. These coordinated attachments resulted communities to fuse or expand, increasing both their number and overall size in NEU stage (Figure 7c).

**Figure 7.**
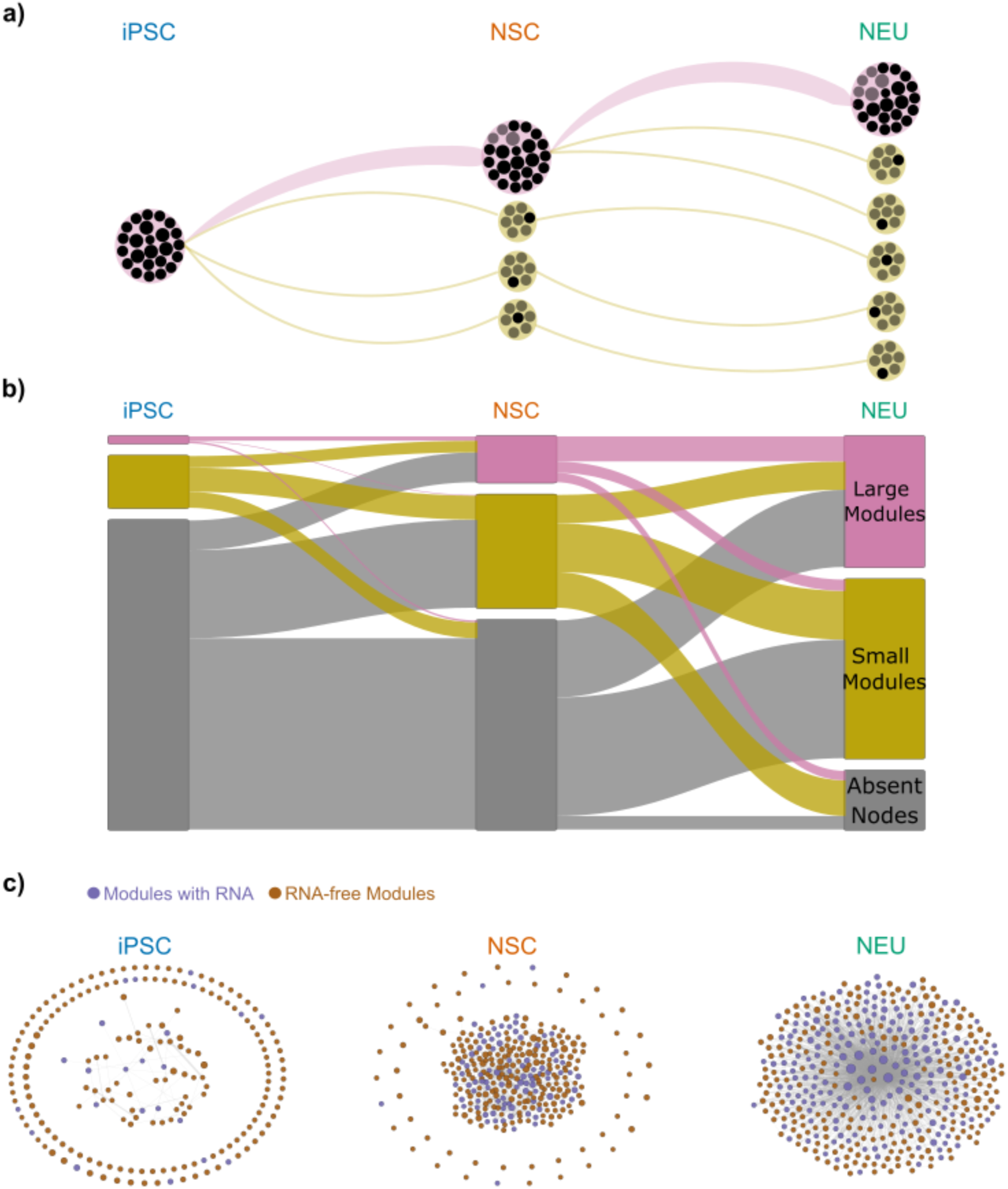
**a)** Schematic of membership changes through division and differentiation. Black nodes (DNA or RNA) are grouped into communities: large (pink; ≥100 nodes) and small (ocher). Dark nodes indicate members of one iPSC community; lighter nodes show mixing with elements from other communities as links are diluted; **b)** Alluvial flow of RNA nodes between Infomap communities across cell types. Large communities are pink; small communities are ocher. Nodes not detected in a given cell type are assigned to an absent block (gray). Block heights encode group sizes and ribbon widths encode transitions; **c)** Snapshots of large-community networks for each cell type. Modules with only DNA–DNA interactions are brown; modules with DNA–DNA plus RNA–DNA interactions are purple. Node size scales with community size. Peripheral “orbit” nodes lack links to other large communities.

We also observed that connectivity increased at two scales. Degree (i.e. the number of connections of a given node) did not have any influence the likelihood of belonging to a large community (β = 2.67×10⁻⁵, *p* = 0.24), meaning that large communities are not simply aggregates of nodes with high number of connections. Within communities, mean degree and clustering strengthened upon differentiation, especially inside large communities: mean degree (i.e. number of contacts) increased from 2.37 (95% CI 2.24–2.51) in iPSC to 3.71 (3.46–3.97) in NSC and 12.66 (10.53–14.80) in NEU; mean clustering rises from 0.068 (0.065–0.071) to 0.200 (0.196–0.205) and 0.401 (0.396–0.406) (Fig. 7c). At the whole-network scale, mean degree also increased, 3.41 (2.78–4.03) in iPSC, 5.13 (3.82–6.44) in NSC, and 10.41 (8.61–12.21) in NEU, accompanying the expansion in the number and average size of large communities (Fig. 7c; Supplementary Figure 7).

Genes that change communities across cell types were more likely to be differentially expressed (1.21-fold, Fisher’s exact test p = 5.74e-6). Moreover, genes that belong to large communities were enriched for transcription factors (2.1-fold, padj = 6e-4, Supplementary Figure 8). Together, these observations indicate that RNA associated interactions contribute to DNA-DNA connectivity in a non-random fashion: RNA seems to not only increases DNA–DNA connectivity within communities but also introduces selective links that draw additional nodes sometimes even entire node groups into large communities as differentiation progressed.

## Discussion

In this study, we integrated high-resolution chromatin interaction maps (HiCap) with genome-wide RNA–chromatin association data (RADICL-seq) across human neural differentiation, providing new insights into how chromatin-associated RNAs may influence cis-regulatory architecture. By bridging DNA-DNA interactomes and RNA–DNA associations, we uncovered a pervasive and dynamic overlap that strongly suggests RNA molecules may act as modulators of enhancer-promoter connectivity, which is also supported by previous studies (*14*, *45–47*). A key finding is that both promoters and enhancers engaged in DNA–DNA interactions were markedly enriched for RNA association, consistent with previous observations that regulatory elements are hotspots for RNA association (*48*). We observed more-than two-thirds of promoters changed their RNA connectivity across neural fate commitment, and these changes influenced transcriptional output and/or cis-connectivity. Based on these findings, we suggest that chromatin-RNA interactions can be predictive of regulatory changes rather than coincidental contacts with the DNA.

We observed an asymmetric role for the RNA associations at promoters versus enhancers. Specifically, RNA dynamic at enhancers strongly coincided with interaction gain or loss. PIR RNA gain was the most robust predictor of interaction gain and similarly loss of RNA at PIRs were consistently overlapped with interaction loss. These observations suggest that enhancer bound RNAs might exert a more dominant influence on contact dynamic and it also aligns with earlier studies where specific lncRNAs acting as molecular tethers or architectural factors modulating local chromatin accessibility or cofactor recruitment (*18*, *45*).

Several well-characterized lncRNAs, including MALAT1, NEAT1, and SNHG14, showed cell type–specific rewiring of their chromatin contacts. When we investigated the direction of regulation of genes whose promoters/enhancers bound to specific RNAs; for a subset of RNAs, their target genes were up or down regulated suggesting additional factors in play in a context dependent manner. However, for some RNAs, the target genes were either up or down-regulated pointing to a systematic effect by the RNA. This dual pattern also supports recent findings that lincRNAs could act as scaffolds for either activating or repressive complexes (*49–51*) and underscores the importance of cellular context in determining RNA function.

We also find that promoters with dynamic RNA associations were more likely to undergo transcriptional changes and concordantly were enriched for neural differentiation–related functions and disease-linked phenotypes, suggesting an active role for chromatin-associated RNAs during the establishment of lineage-specific gene expression programs.

We coarse-grained the interaction data into large modules, or *communities*, which are groups of nodes that interact more frequently with each other than with the rest of the network. Across differentiation, we observed that nodes often shifted communities, indicating a dynamic reorganization of the higher-order chromatin structure. These changes were not dominated by individual hubs and often involved movement of groups of nodes, frequently linked through RNA-mediated contacts (DNA-RNA-DNA). This supports a model in which caRNAs help connect or stabilize subsets of regulatory elements.

Our integrative approach highlights methodological advantages. By combining HiCap with RADICL-seq, we overcome the limitations of each assay in isolation: HiCap provides precise promoter–enhancer maps but lacks information on RNA associations, while RADICL-seq captures RNA binding but not 3D chromatin topology. Nevertheless, our study has several limitations. First, RADICL-seq data does not distinguish between direct RNA–DNA base-pairing and indirect interactions mediated by RNA-binding proteins. Experimental data using orthogonal approaches such as CHART (*52*), ChIRP (*53*), or CRISPR-based RNA tethering (*54*) would provide insight the mechanistic nature of these associations. Second, our experiments were done on steady-state cell populations, however, using single-cell or time-resolved data could help to refine the temporal dynamics of chromatin-RNA binding effects on the interactome. Third, we cannot distinguish whether associated RNAs at different interacting DNA loci involves the same individual RNA or multiple copies of the same RNA species bound simultaneously at distinct regions. Therefore, although at a network level we can infer co-location of RNA and DNA regions, we cannot resolve whether one RNA molecule or many copies of the same RNA underlie these multi-locus associations. Finally, although our results strongly implicate RNAs as regulators of chromatin interactions, functional perturbation of candidate RNAs is necessary to establish causality.

In summary, our study provides a comprehensive view of how chromatin-associated RNAs overlap with promoter–enhancer networks to shape transcriptional programs in neural differentiation. We believe our work will help to advance our understanding of chromatin-associated RNAs and their role in shaping genome regulation and structure.

## Notes

https://kth-my.sharepoint.com/:f:/g/personal/pelinak_ug_kth_se/IgDz7JCWB0VJSJtzmosi0luPAfvlWdhNgRCPAKQIz6soLkU?e=C84z3o

### Competing Interest Statement

The authors have declared no competing interest.

### Funding

- European Commission: 860002

