## Supplementary Figures for "Extending cis-regulatory networks using chromatin-RNA interactions"

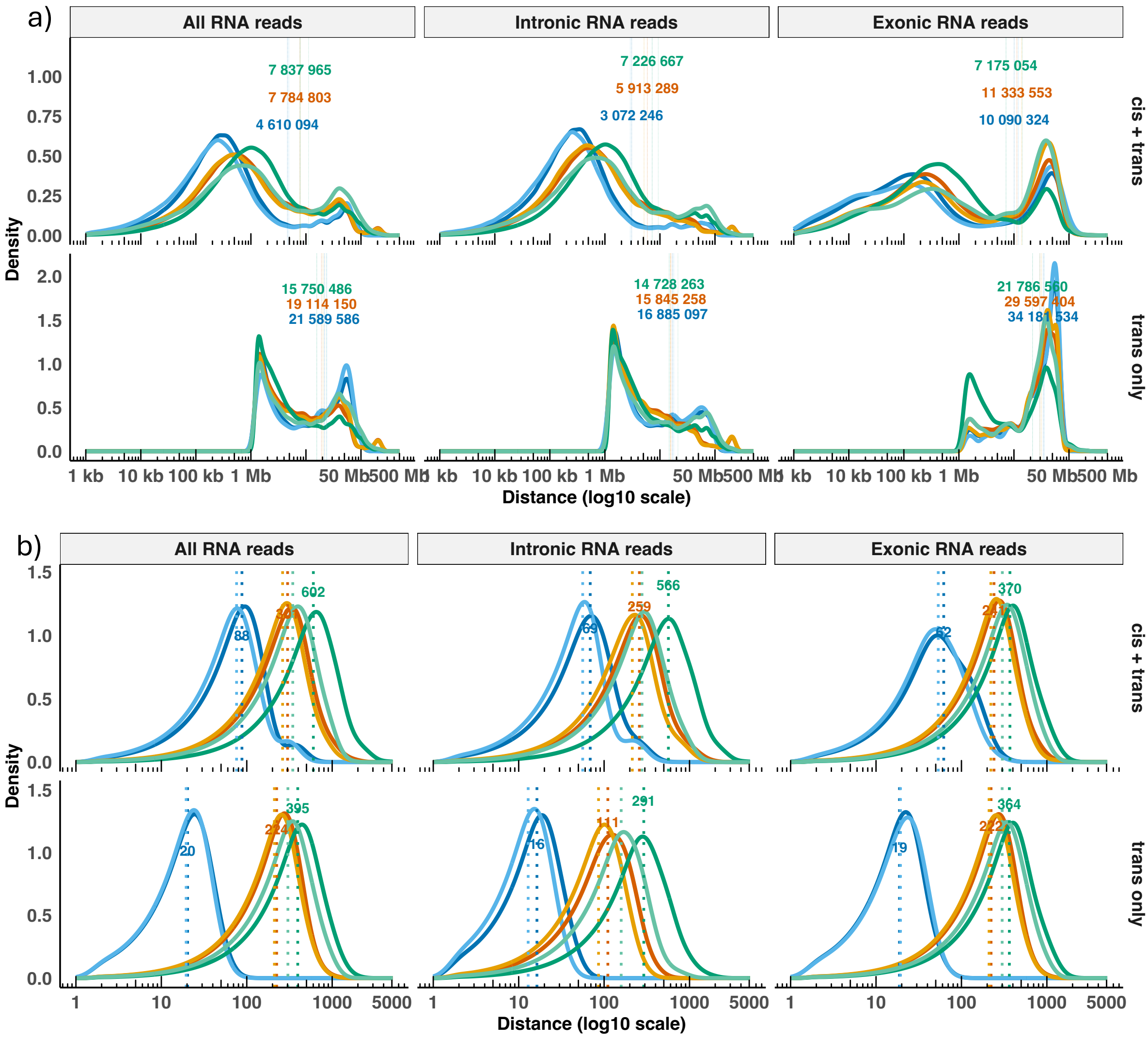


***Supplementary Figure 1***

*The average a) width of the DNA regions b) weight of interactions in RADICL-seq dataset stratified by edge type and distance*

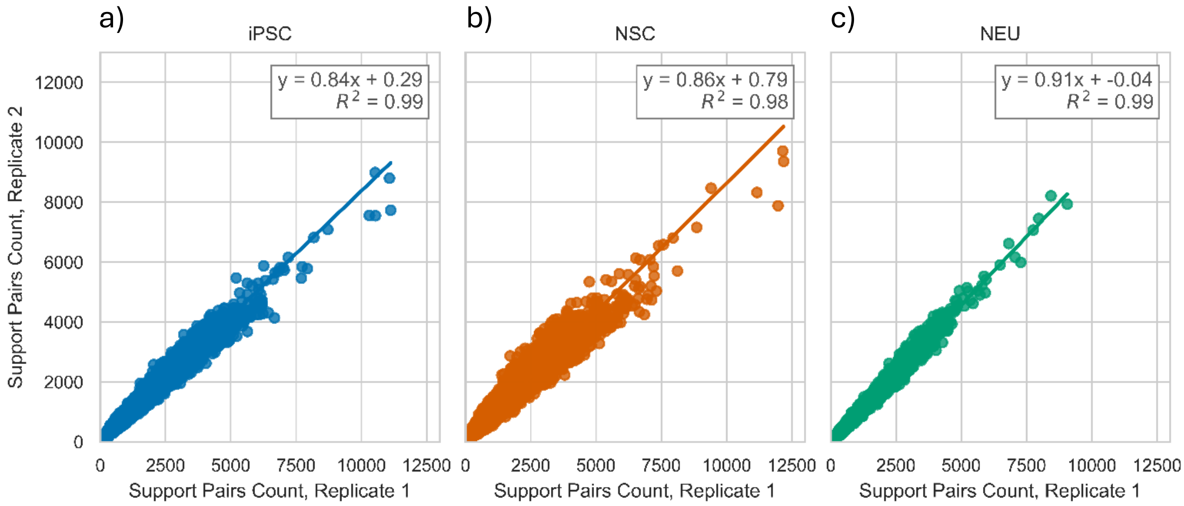


d

| Cell Type | Source RNA | Target DNA | Number of Interactions |
|---|---|---|---|
| iPSC.rep1 | 13 384 | 395 573 | 600 391 |
| iPSC.rep2 | 10 327 | 264 826 | 357 011 |
| NSC.rep1 | 14 303 | 649 765 | 1 056 048 |
| NSC.rep2 | 13 533 | 701 705 | 1 019 522 |
| NEU.rep1 | 17 992 | 575 970 | 1 344 701 |
| NEU.rep2 | 10 822 | 547 411 | 934 621 |

e)

| Interaction | iPSC | NSC | NEU |
|---|---|---|---|
| RNA - Prom\|RNA(+) | 14 036 ± 2 937 | 42 500 ± 4 140 | 83 940 ± 13 972 |
| RNA - GWAS\|RNA(+) | 535 ± 101 | 1 903 ± 147 | 4 552 ± 718 |
| RNA - PIR\|RNA(+) | 79 980 ± 18 930 | 320 445 ± 29 189 | 757 006 ± 131 667 |
| Prom\|RNA(-) - Prom\|RNA(-) | 7 402 ± 733 | 3 580 ± 59 | 4 822 ± 84 |
| Prom\|RNA(-) - Prom\|RNA(+) | 7 504 ± 414 | 8 621 ± 98 | 12 250 ± 204 |
| Prom\|RNA(-) - GWAS\|RNA(-) | 268 ± 19 | 156 ± 1 | 165 ± 8 |
| Prom\|RNA(-) - GWAS\|RNA(+) | 133 ± 12 | 168 ± 1 | 317 ± 6 |
| Prom\|RNA(-) - PIR\|RNA(-) | 44 660 ± 6 754 | 18 635 ± 682 | 25 467 ± 980 |
| Prom\|RNA(-) - PIR\|RNA(+) | 18 327 ± 2 093 | 27 672 ± 1 388 | 29 587 ± 1 510 |
| Prom\|RNA(+) - Prom\|RNA(+) | 2 421 ± 390 | 5 514 ± 92 | 8 432 ± 147 |
| Prom\|RNA(+) - GWAS\|RNA(-) | 206 ± 18 | 182 ± 5 | 248 ± 8 |
| Prom\|RNA(+) - GWAS\|RNA(+) | 120 ± 22 | 352 ± 14 | 514 ± 11 |
| Prom\|RNA(+) - PIR\|RNA(-) | 27 737 ± 3 150 | 30 820 ± 1 897 | 33 648 ± 1 669 |
| Prom\|RNA(+) - PIR\|RNA(+) | 17 686 ± 3 478 | 63 883 ± 3 244 | 79 836 ± 3 582 |
| GWAS\|RNA(-) - GWAS\|RNA(-) | 21 ± 3 | 10 ± 1 | 9 ± 1 |
| GWAS\|RNA(-) - GWAS\|RNA(+) | 10 ± 2 | 16 ± 2 | 17 ± 0 |
| GWAS\|RNA(-) - PIR\|RNA(-) | 241 ± 37 | 96 ± 12 | 116 ± 3 |
| GWAS\|RNA(-) - PIR\|RNA(+) | 94 ± 12 | 119 ± 8 | 116 ± 12 |
| GWAS\|RNA(+) - GWAS\|RNA(+) | 3 ± 1 | 11 ± 2 | 25 ± 2 |
| GWAS\|RNA(+) - PIR\|RNA(-) | 64 ± 17 | 98 ± 4 | 104 ± 3 |
| GWAS\|RNA(+) - PIR\|RNA(+) | 43 ± 15 | 236 ± 8 | 284 ± 3 |

f)

| Interaction | iPSC | NSC | NEU |
|---|---|---|---|
| RNA - Prom\|RNA(+) | 7 045 | 31 093 | 49 284 |
| RNA - GWAS\|RNA(+) | 222 | 1 252 | 2 479 |
| RNA - PIR\|RNA(+) | 24 760 | 160 570 | 336 201 |
| Prom\|RNA(-) - Prom\|RNA(-) | 4 080 | 2 412 | 3 898 |
| Prom\|RNA(-) - Prom\|RNA(+) | 3 790 | 5 806 | 9 873 |
| Prom\|RNA(-) - GWAS\|RNA(-) | 135 | 106 | 126 |
| Prom\|RNA(-) - GWAS\|RNA(+) | 59 | 95 | 256 |
| Prom\|RNA(-) - PIR\|RNA(-) | 18 443 | 10 640 | 17 592 |
| Prom\|RNA(-) - PIR\|RNA(+) | 6 408 | 15 581 | 18 972 |
| Prom\|RNA(+) - Prom\|RNA(+) | 1 275 | 3 783 | 6 913 |
| Prom\|RNA(+) - GWAS\|RNA(-) | 98 | 119 | 196 |
| Prom\|RNA(+) - GWAS\|RNA(+) | 56 | 251 | 422 |
| Prom\|RNA(+) - PIR\|RNA(-) | 9 860 | 17 555 | 21 735 |
| Prom\|RNA(+) - PIR\|RNA(+) | 6 976 | 38 782 | 54 553 |
| GWAS\|RNA(-) - GWAS\|RNA(-) | 14 | 8 | 7 |
| GWAS\|RNA(-) - GWAS\|RNA(+) | 7 | 7 | 12 |
| GWAS\|RNA(-) - PIR\|RNA(-) | 97 | 55 | 76 |
| GWAS\|RNA(-) - PIR\|RNA(+) | 29 | 62 | 63 |
| GWAS\|RNA(+) - GWAS\|RNA(+) | 1 | 6 | 21 |
| GWAS\|RNA(+) - PIR\|RNA(-) | 12 | 52 | 65 |
| GWAS\|RNA(+) - PIR\|RNA(+) | 14 | 139 | 173 |

***Supplementary Figure 2***

*a-c) Scatter plots of supporting pairs of replicate 1 versus replicate 2 for each cell type, d) statistically significant RNA-DNA proximities for each cell type and replicate, e) The combined number of HiCap / RADICL-seq interactions in all networks, grouped by cell type and averaged over all replicates. f) The number of HiCap / RADICL-seq interactions in the networks present in all replicates, grouped by cell type.*

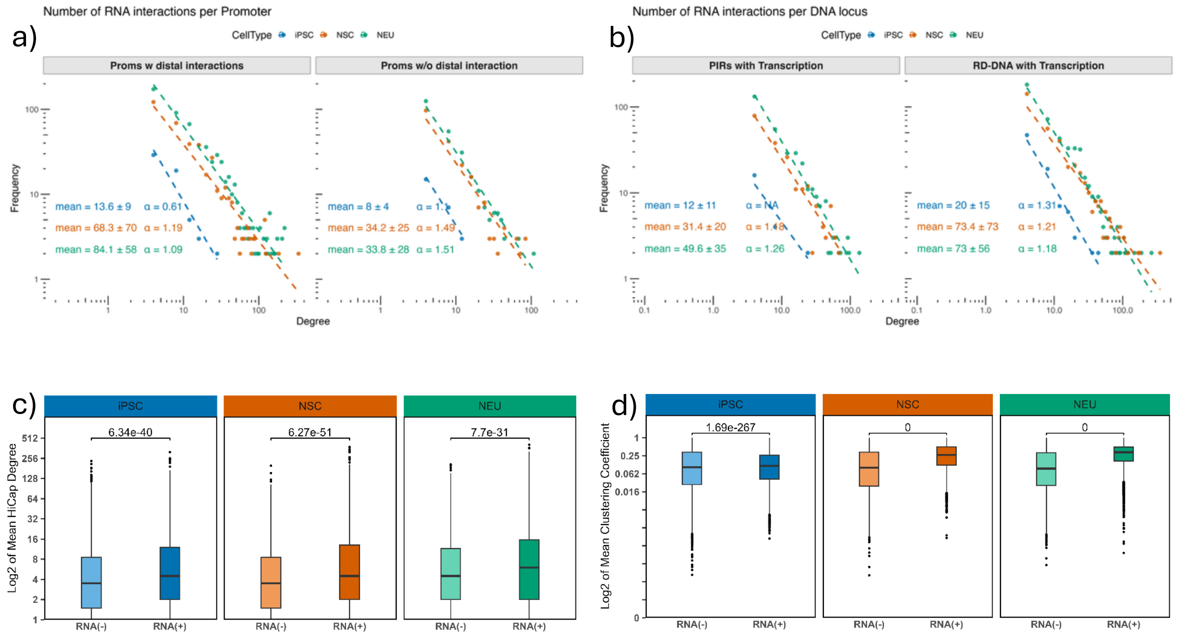

***Supplementary Figure 3***

*a) RNA contact degree distribution of HiCap promoters and promoters with no distal interaction, x-axis denotes the degree (the number of interacting RNA transcripts) of promoters, y-axis corresponds to the number of promoters with that degree. The slope of the line fits are shown on the top right b) The RNA contact degree distribution of HiCap interacting regions (PIRs, excluding promoters) and DNA regions (excluding all promoters and PIRs). Only trans RNA-DNA interactions are used. The c) HiCap degree and d) clustering coefficient of promoters with respect to their RNA interaction status, considering only trans RNA-DNA interactions.*

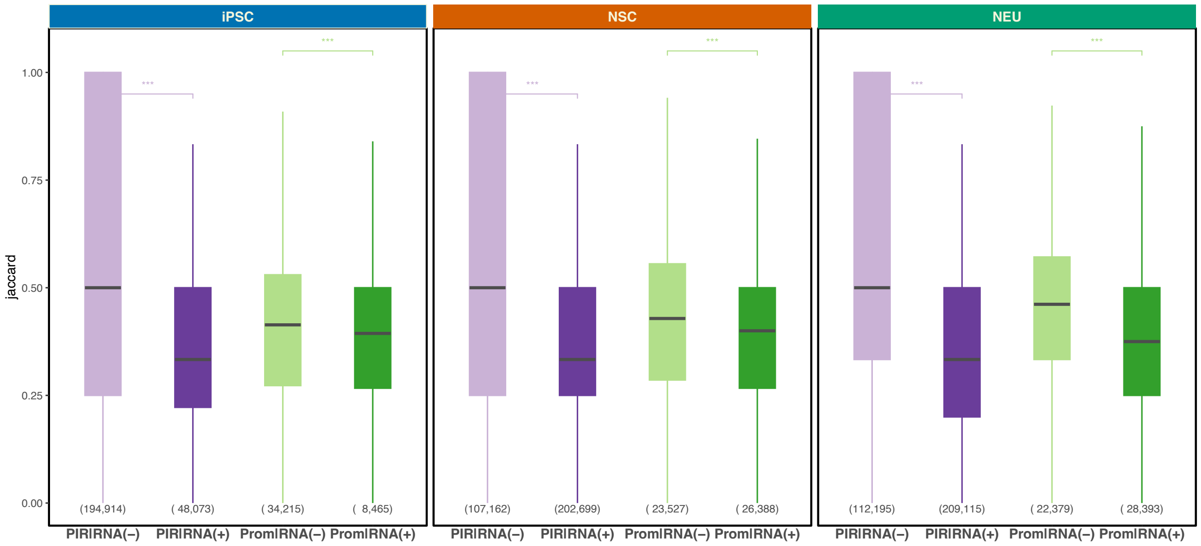


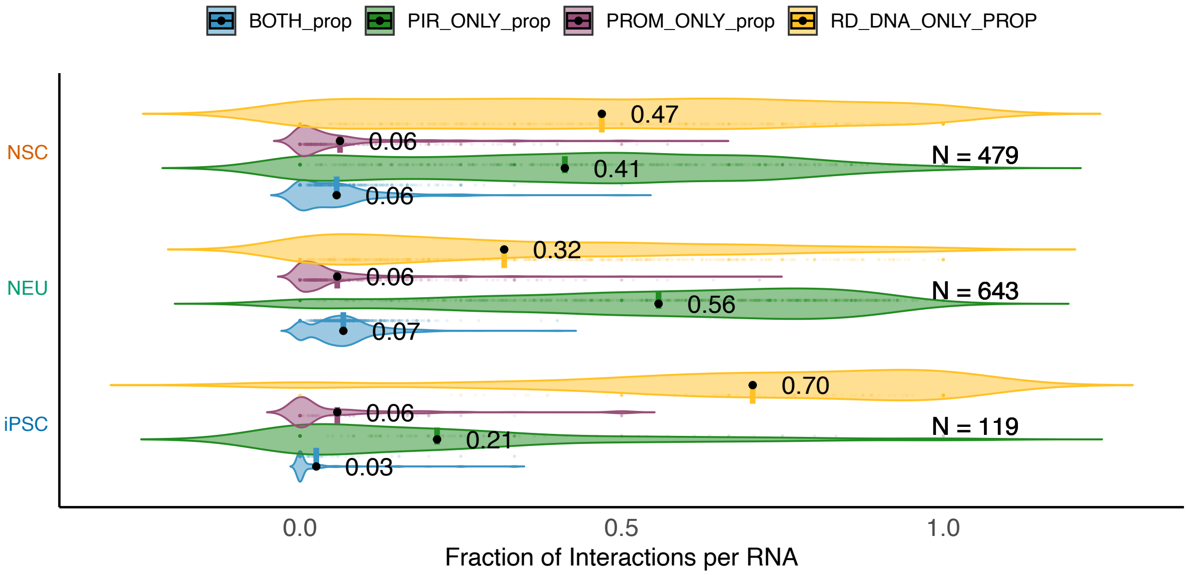


***Supplementary Figure 4.***

*a) The jaccard distribution of interacting DNA elements (PIRs and promoters) with respect to their RNA interaction status. b) For each RNA, we counted number of Promoter, PIR and RD-DNA (DNA with no HiCap overlap) and divided to the total number of interactions. The x-axis shows the distribution of such proportions for all interacting RNAs. Only trans RNA-DNA interactions are taken into account. c)*

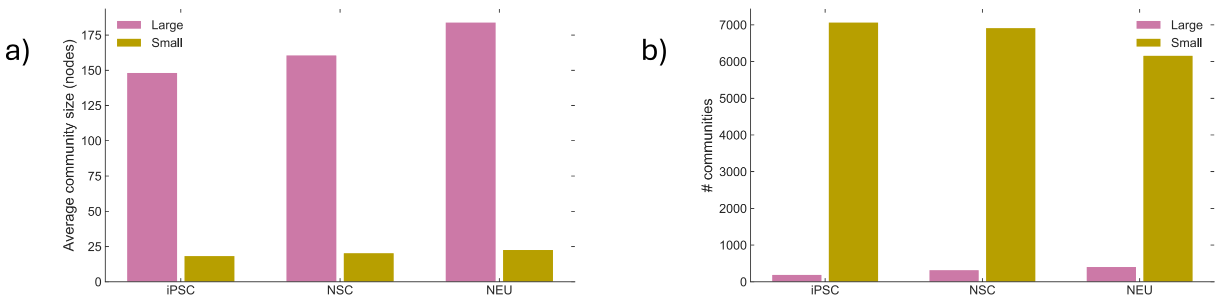


***Supplementary Figure 5.***

*Community size and counts across differentiation. a) Mean community size for large (≥100 nodes) and small (<100 nodes) communities in iPSC, NSC, and NEU. b) Number of communities by category and cell type: large communities and small.*

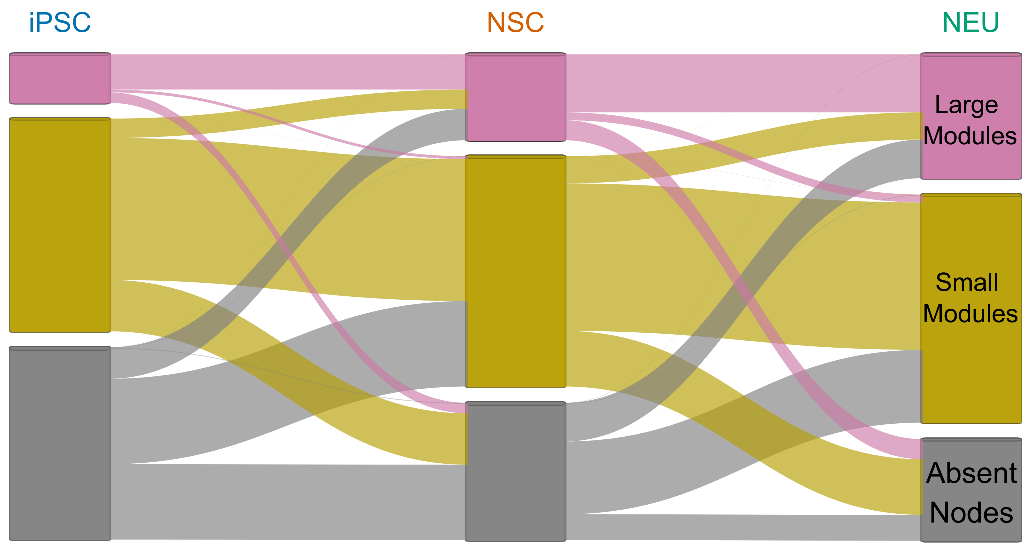


***Supplementary Figure 6.***

*Alluvial flow of all nodes across differentiation. Alluvial diagram showing community-category flows for all nodes (DNA and RNA) from iPSC to NSC to NEU. Blocks encode community categories, large (pink; ≥100 nodes), small (ocher; <100 nodes), and absent (gray, not detected in that cell type). Block heights indicate counts per category; ribbon widths show transitions between categories. Compare with the RNA-only alluvial in Fig. 7b.*

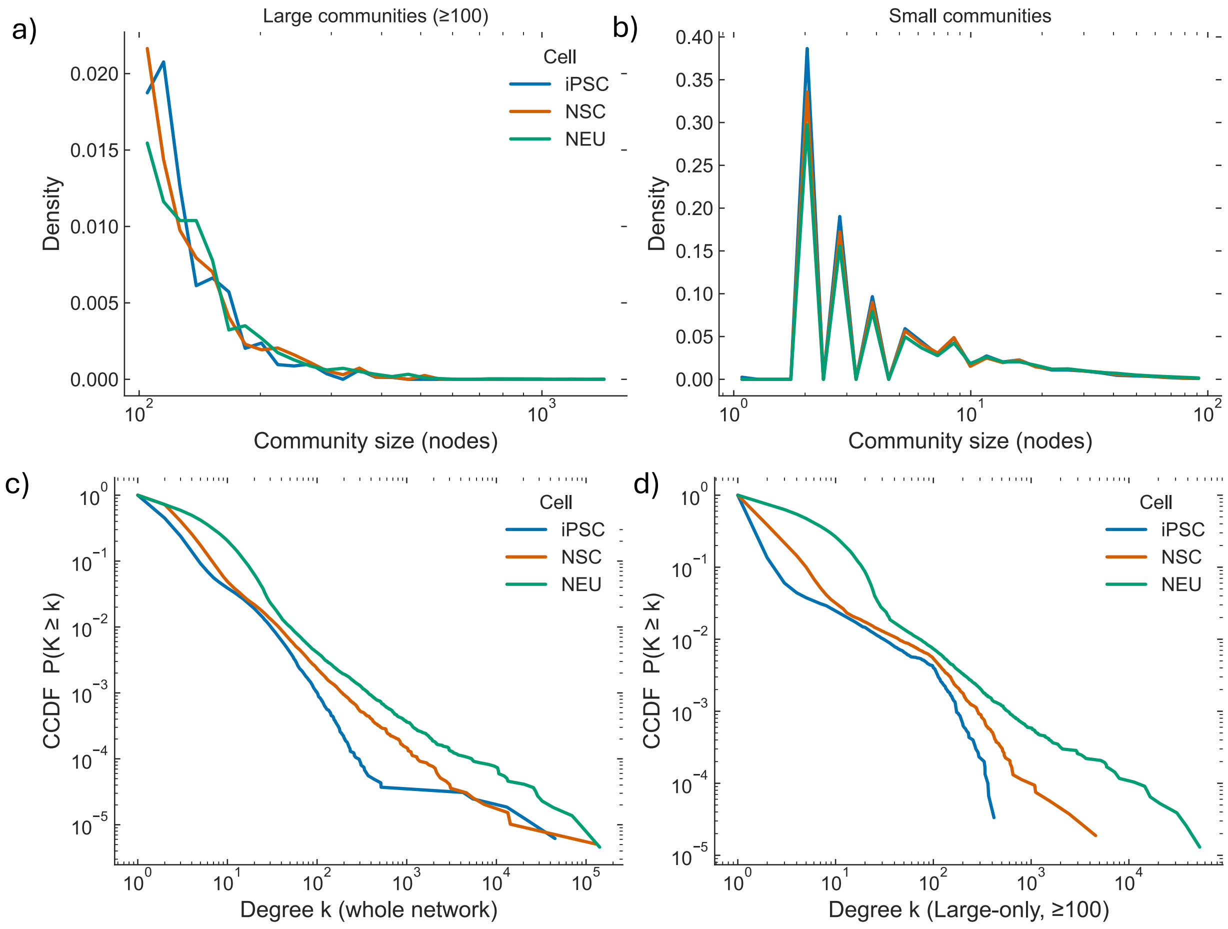


***Supplementary Figure 7.***

*Community size and degree distributions across differentiation. a) Size distribution of large communities (≥100 nodes) for iPSC, NSC, and NEU. b) Size distribution of small communities (<100 nodes). c) Complementary cumulative distribution function (CCDF) of node degree K for nodes within large communities; CCDF defined as P(K' ≥ K). d) CCDF of node degree K for nodes within small communities. Counts by cell type: large communities = 202 (iPSC), 331 (NSC), 418 (NEU); small communities = 7,077, 6,924, 6,171, respectively.*

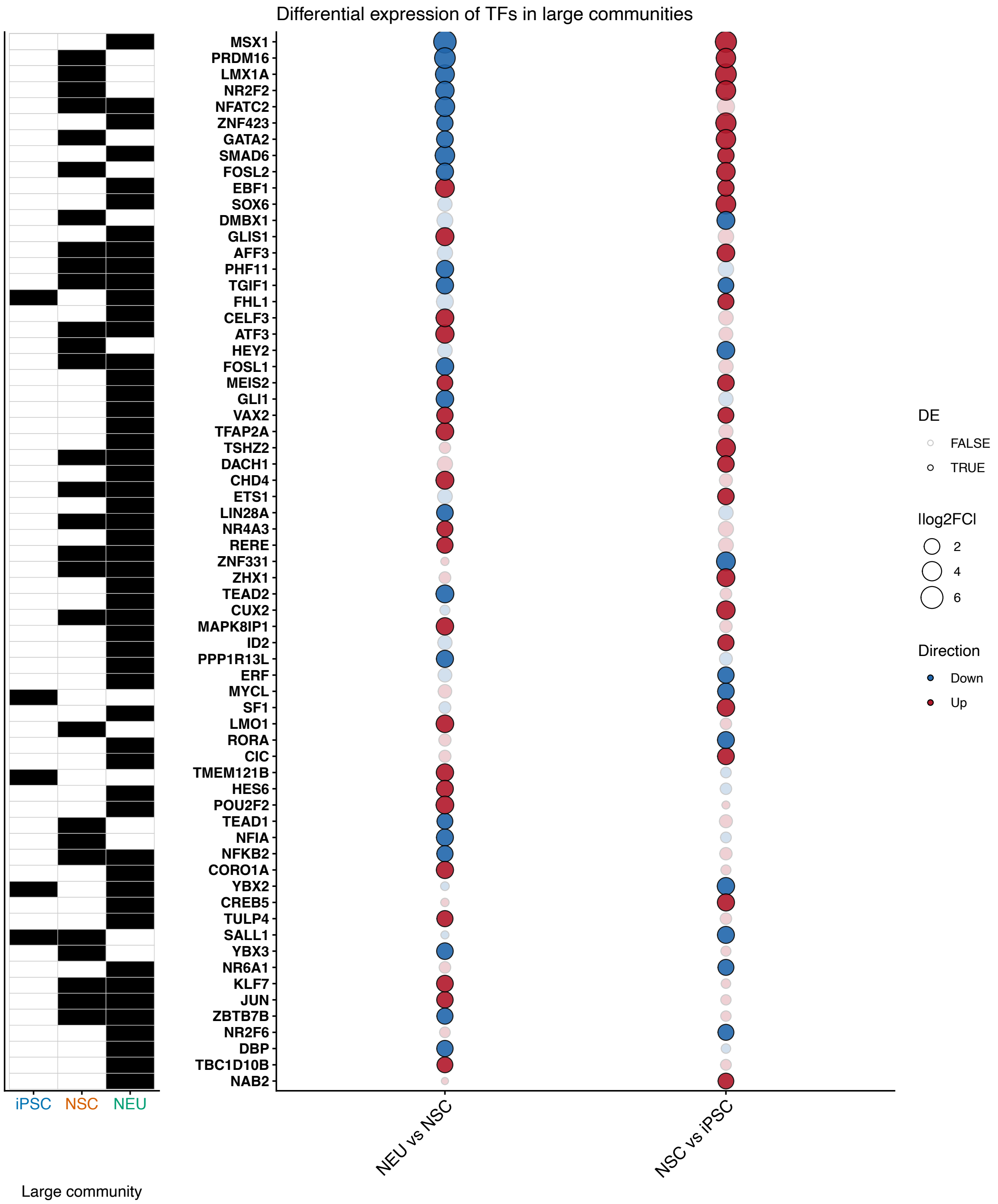


***Supplementary Figure 8.*** *The transcription factors that belong to a large community and differentially expressed in at least one cell type and their log2-fold expressed change across the consecutive cell types.*
